# Maternal Immune Activation Disrupts Epigenomic and Functional Maturation of Cortical Excitatory Neurons

**DOI:** 10.1101/2025.04.28.651094

**Authors:** Chi-Yu Lai, Jessica Arzavala, Antonio Pinto-Duarte, Shiyuan Wang, Junhao Li, Hanqing Liu, Julia Osteen, Rosa Gomez Castanon, Joseph Nery, Susan B. Powell, Joseph R. Ecker, Eran A. Mukamel, M. Margarita Behrens

## Abstract

Elevated levels of maternal pro-inflammatory cytokines during gestation can disrupt offspring neural development, increasing the risk of neurodevelopmental disorders. We studied the effects of Poly(I:C)-induced maternal immune activation (PIC-MIA) during mid-gestation on developing cortical excitatory neurons’ DNA methylation and transcriptome. PIC-MIA disrupted the developmental regulation of synapse-related genes and of genes implicated in autism spectrum disorders. Genomic regions that gain or lose DNA methylation during normal development were altered following PIC-MIA, including neurodevelopmental transcription factor binding sites. The DNA methylation and transcriptional changes were consistent with a delay in excitatory neuron maturation. Whole-cell recordings showed that PIC-MIA preferentially altered the physiological development of layer 5 excitatory neurons. Taken together, present results suggest that alterations in the epigenome, through the disruption of circuit formation, may drive the long-term consequences of maternal infection during gestation.

## Introduction

Epidemiological and experimental evidence suggests early life adversity, such as disruption of the maternal environment during prenatal development, is a common factor in the etiology of autism spectrum disorder (ASD) and schizophrenia.^1,2^ In rodents, maternal immune activation (MIA) during early- to mid-embryonic development leads to permanent neurodevelopmental alterations in the offspring,^3–6^ including behavioral phenotypes that resemble those of human autism spectrum disorder (ASD) such as social impairments and behavioral inflexibility.^7–9^

The neurodevelopmental alterations observed in MIA offspring could be caused by dysregulation of the brain epigenome,^10–13^ and ensuing altered gene expression.^14–16^ We previously showed that DNA methylation in the frontal cortex during embryonic and early postnatal development is highly dynamic^17^ and marked by the accumulation of non-CG methylation at the same time as the period of synaptogenesis in mice and humans. The dynamic changes in brain DNA methylation patterns between gestation and early postnatal life^17^ suggest the epigenome could be particularly vulnerable to environmental disruption during this period.

How the activation of maternal immune response and the offspring’s epigenetic program interact to influence brain gene expression and trigger behavioral phenotypes remains largely unknown. Studies of cortical gene expression in the adult offspring of MIA, induced by treatment of pregnant dams with the viral mimetic Polyriboinosinic–polyribocytidylic acid (Poly(I:C)) (PIC-MIA), have reported relatively subtle alterations.^8,11,12,18^ This is also true when the embryos are analyzed a few hours after MIA in rats or mice.^14,19–22^ Recent data showed that PIC-MIA offspring developed transcriptional dysregulation in cortex several days after the maternal immune activation occurred, which was accompanied by alterations in corticogenesis including alterations in precursor cell proliferation and early neuronal maturation marker expression.^22^ These effects could be caused by disruption of the normal development of the epigenome of specific brain cell types. We previously showed neuronal DNA methylation patterns are highly dynamic during the perinatal period, spanning birth to adolescence when adult patterns are established.^17^ We further showed adult DNA methylation and transcriptional patterns are cell-type specific.^23–25^ However, little is known about when these patterns are established in each cell type and whether changes in the maternal environment influence this epigenomic program.

Our previous work showed a correlation between frontal cortex transcriptional alterations and disrupted cognitive flexibility in adult mice,^8^ a task dependent on frontal cortex function.^26–28^ In this study we focused specifically on the excitatory class of cortical neurons during development and tested the hypothesis that PIC-MIA alters the development of these neurons through disruption of the epigenome. We used whole-genome DNA methylome and transcriptome analysis to examine how maternal immune activation with Poly(I:C) perturbs the epigenomic developmental dynamics of frontal cortex excitatory neurons during the perinatal period. We found that PIC-MIA alters DNA methylation and transcription during the perinatal period, long after the maternal immune activation has subsided, with consequences for the physiology of specific excitatory neuron cell types. Our findings highlight how maternal immune activation (PIC-MIA) significantly delays DNA methylation patterns and disrupts physiological maturation of cortical excitatory neurons, potentially contributing to behavioral abnormalities observed in neurodevelopmental disorders. These results add novel insight into the epigenetic mechanisms through which prenatal immune challenges influence neurodevelopment, particularly emphasizing lasting impacts on cortical circuitry.

## Results

### Developmental DNA methylation and transcriptional changes in excitatory neurons during the perinatal period

DNA methylation in neurons undergoes a profound reconfiguration during postnatal development in mice and humans.^17^ Mature neurons have cell-type-specific patterns of DNA methylation that reflect the specialized regulation of excitatory and inhibitory neuron subtypes.^17,23,24,29,30^ We analyzed the development of frontal cortex excitatory neurons at four key ages: embryonic day 15 (E15), and postnatal days P0, P13, and P39 (Fig. 1a). We used the INTACT mouse line^23^ (on a C57BL/6J background) crossed to a Cre-expressing line (Neurod6-Cre, or Nex-Cre)^31,32^ to target excitatory neuronal nuclei early during embryonic development.^33^

**Fig. 1.**
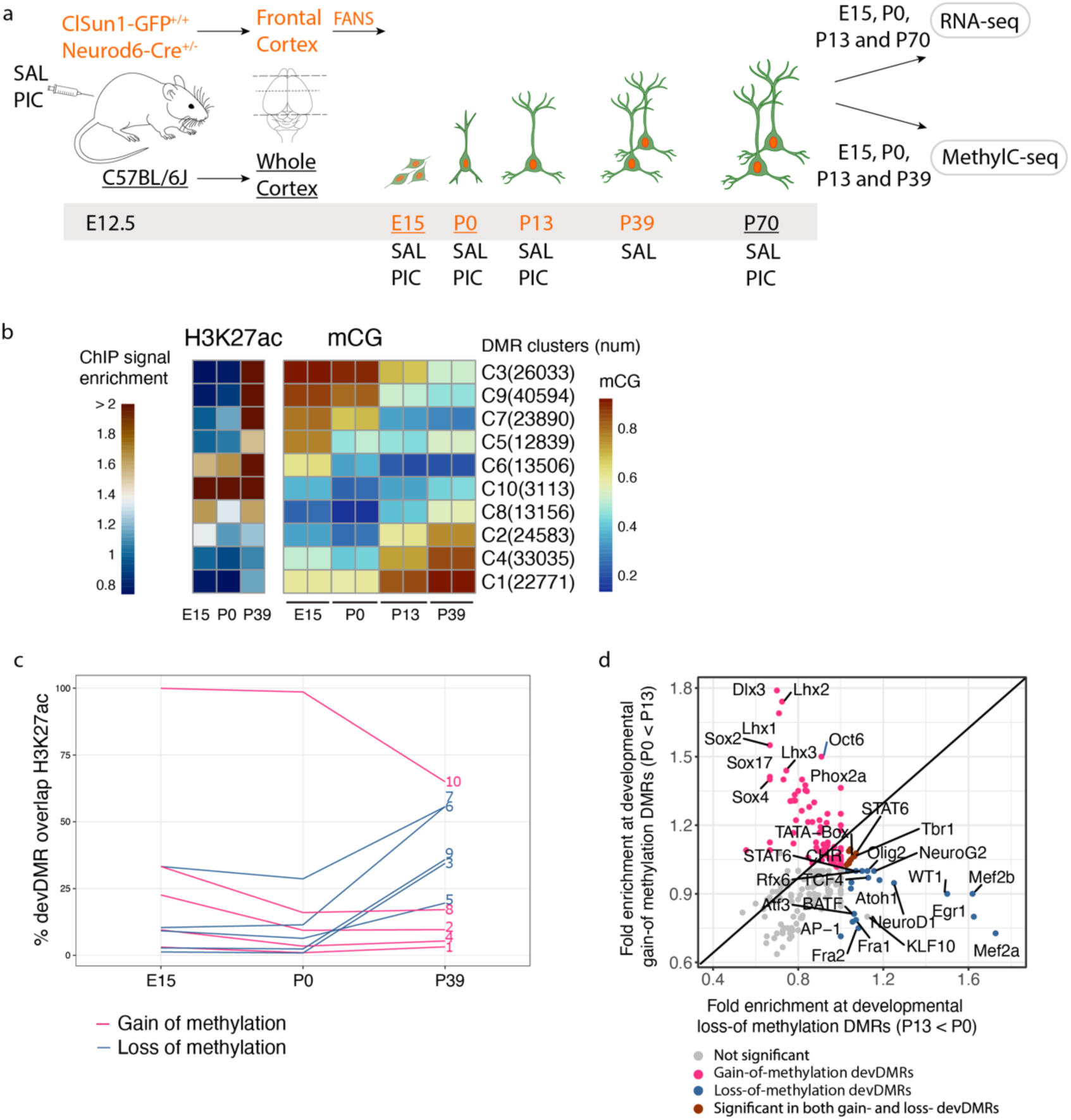
DNA methylation dynamics in frontal cortex excitatory neurons during pre- and postnatal brain development. **a.** Schematic diagram of the study design. PIC, Poly(I:C) treatment. SAL, saline treatment. E12.5, embryonic day 12.5. E15, embryonic day 15. P0, postnatal day 0. P13, postnatal day 13. P39, postnatal day 39. P70, postnatalday 70. FANS, fluorescence-activated nuclei sorting. Samples that underwent fluorescence-activated nuclei sorting (FANS) are colored in orange. Underlined whole-cortex time points (E15, P0, and P70) only underwent RNA-seq. **b.** Mouse excitatory neurons of cortex developmental DMRs (devDMRs). Differentially methylated regions (DMRs) were identified during mouse brain development with samples of the cortex (E15) and fluorescence-activated cell-sorted nuclei of excitatory neurons (P0, P13, and P39). Pairwise differentially methylated regions (≥30% methylation difference, p<1×10-5) at CG dinucleotides were identified using DSS. Groups of DMRs with similar patterns of average methylation level (mCG/CG) across samples were clustered using k-means clustering (k=10). The resulting clusters were arranged to show developmental gain-of-methylation (clusters 1, 2, 4, 8, and 10) and loss-of-methylation (clusters 3, 5, 6, 7, and 9). The heatmap shows the DNA methylation level in the CG context (mCG) of DMRs. H3K27 acetylation (H3K27ac) ChIP-Seq data for forebrain development at E15,^34^ P0 and P39 were used to plot the mean H3K27ac signal (enrichment compared to input DNA) in these DMR clusters. **c.** The percentage of devDMRs overlapping H3K27ac peaks at E15, P0, and P39. This data suggests a dynamic change in active enhancers during the developmental period, with the great majority of embryonic active enhancers being silenced (gain-of-methylation; cluster 1, 2, 4,8, and 10; pink lines) as development progresses, while mature neuron enhancers demethylate (loss-of-methylation; clusters 3, 5, 6, 7 and 9 blue lines) during the postnatal period. **d.** Enriched transcription factor binding motifs in devDMRs. Pink/blue dots are significantly enriched transcription factor binding motifs in developmental gain-of-methylation/loss-of-methylation from P0 to P13.

Excitatory neuron development profoundly changes DNA methylation at CG dinucleotides (mCG) between embryonic day 15 (E15) and 6 weeks of postnatal age (P39) (Fig. 1b, Supplementary Table 1), as previously shown at the level of whole frontal cortex tissue.^17^ We identified 213,521 developmental differentially methylated regions (devDMRs) that gain or lose mCG between E15 and P39. The pattern of DNA methylation dynamics was diverse, with thousands of devDMRs gaining or losing methylation between each sequential stage of development from prenatal (E15) to birth (P0), early development (P13) and adulthood (P39). The largest number of devDMRs corresponded to early waves of DNA methylation gain or loss during the first two postnatal weeks (Fig. 1b, clusters 1, 2 and 4 gaining methylation, and clusters 3, 7 and 9 losing methylation).

DNA methylation works together with other epigenetic modifications, such as histone modifications associated with active enhancers and promoters (H3K27 acetylation), that have their own dynamic patterns of regulation during forebrain development.^34^ To address the relationship between these epigenetic marks, we compared our data with H3K27ac chromatin immunoprecipitation followed by sequencing (ChIP-seq) data from mouse forebrain at E15, P0,^34^ and at P39^33^ (Supplementary Table 2). We found that devDMRs which gain methylation (96,658 devDMRs, clusters 1, 2, 4, 8, and 10 in Fig. 1b) overlapped more with active regulatory regions (H3K27ac peaks) at E15 (17.5%) than at P0 (9.2%) or P39 (9.4%). This is consistent with a wave of silencing of putative enhancer regions during the transition between E15 and birth. On the other hand, loss-of-methylation devDMRs (116,862 regions, clusters 3, 5, 6, 7, and 9 in Fig. 1b) overlapped relatively few H3K27ac peaks at E15 (8.2%) and P0 (7.4%) but substantially increased their activity at P39 (40.1%) (Fig. 1c). These regions represent a postnatal wave of activation of putative-enhancer regions.

To address the regulatory significance of devDMRs, we analyzed the enrichment of DNA binding motifs for specific transcription factor (TF) families within devDMRs (Fig. 1d). We found that devDMRs that gain methylation overlap TF binding motifs of early-development transcription binding sites, including *Sox* and *Lhx*. By contrast, devDMRs that lose mCG during development were enriched in motifs of TFs involved in neuronal maturation, including *Neurod* and *Neurog* (Supplementary Table 3).

In parallel with the DNA methylation dynamics, gene expression for thousands of genes increased or decreased during the perinatal period (Supplementary Table 4) between E15 and P0 (Fig. 2a, left) and between P0 and P13 (Fig. 2a, center). By P13, transcriptional changes began to stabilize, with fewer genes significantly changing between P13 and P39 (Fig. 2a, right). To better understand these developmental transitions, we examined gene expression patterns of genes involved in excitatory neuron development and function. Using the molecular signatures database,^35,36^ we selected genes that are key to processes in cortical circuit development such as synaptic activity, glutamate receptor signaling, and calcium channel activity. At E15, synaptic genes, including those involved in glutamatergic and GABAergic neurotransmitter signaling in excitatory neurons, had relatively low expression (Fig. 2b, left), consistent with the early stage of neuronal development. In contrast, transcription factors (TFs) critical for excitatory cortical neuron development (*Sox* and *Lhx* families) had higher expression levels at this embryonic stage (Fig. 2b, right), reflecting active regulation of neuronal specification. By P0 and P13, synaptic gene families (genes encoding GABA and glutamate receptor subunits) showed markedly higher expression, reflecting increased synaptic activity and maturation of excitatory circuits.

**Fig. 2.**
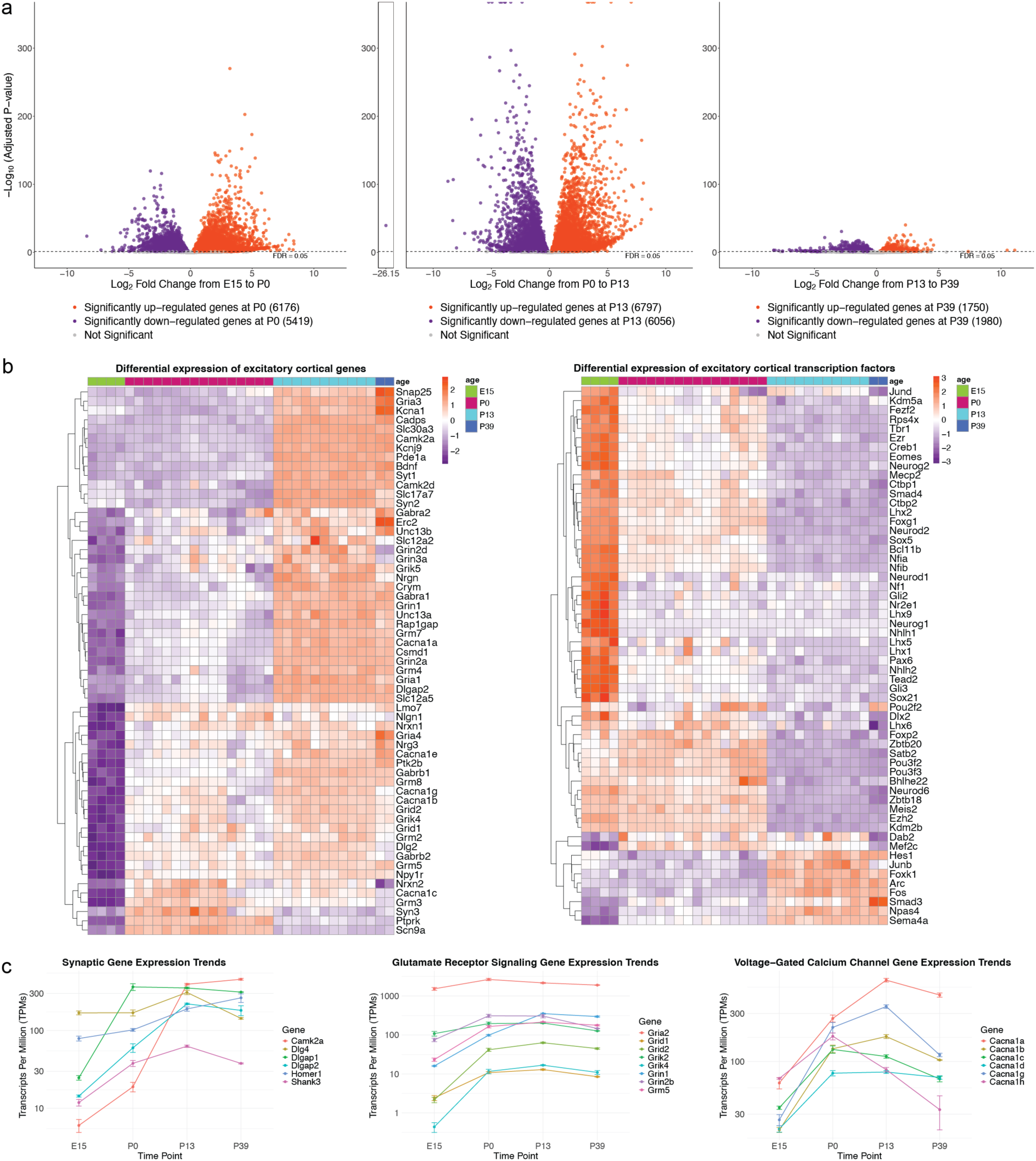
Transcriptional changes during normal cortical excitatory neuron development. **a.** Volcano plots showing differential gene expression in cortex across developmental stages: embryonic day 14 (E14) vs. postnatal day 0 (P0) (left), P0 vs. P13 (center), and P13 vs. P39 (right). Genes up-regulated in the older time point are highlighted in orange, while those down-regulated are shown in purple. Gray points represent genes that are not significantly differentially expressed. **b.** Heatmaps displaying variance-stabilized expression levels of genes (left) and transcription factors (right) specific to cortical excitatory neurons and neurotransmitter receptors. Each gene was re-scaled by Z-score normalization and ordered by hierarchical clustering. Orange indicates higher expression levels while purple denotes lower expression. **c.** Trends in averaged transcripts per million (TPMs) for gene families involved in excitatory cortical development and function.

We further explored the transcriptional changes underlying normal excitatory cortical neuron development using enrichment of biological process functional annotations^37,38^ (Supplementary Fig. 1). Between E15 and P0, up-regulated genes were enriched in synaptic functions, such as *chemical synaptic transmission* and *transmembrane transport* (Supplementary Fig. 1a) and continue to be up-regulated between P0 and P13 (Supplementary Fig. 1b), highlighting the initiation of synapse assembly during the transition from embryonic to postnatal stages. In contrast, by P13 to P39, the enrichment shifted, with down-regulated genes associated with *synapse organization* and *axonogenesis* (Supplementary Fig. 1c). This suggests that synapse formation and circuit assembly are largely complete by P39, underscoring the critical changes in brain structure and function during the perinatal period.

To relate the early postnatal dynamics of excitatory neuron gene expression to cognitive development and function, we examined the expression trends of genes involved in the postsynaptic density in glutamatergic neurons,^39^ glutamate receptors,^40^ and calcium channels.^41^ Between E15 and P0, synaptic genes such as *Camk2a* and *Dlgap2* show an upward trend that stabilizes at P13 and P39 (Fig. 2c, left), consistent with their roles in postsynaptic formation and function. Expression of glutamate receptor-signaling genes and voltage-gated calcium channel genes followed a similar trajectory, peaking around P0 and stabilizing by P13 and P39 (Fig. 2c, center and right). These developmental trajectories provide a reference for understanding the transcriptional changes induced by MIA.

### PIC-MIA disrupts excitatory neuron gene expression of offspring at birth

Gene expression in brain cells of MIA offspring is altered during late gestation following Poly(I:C) exposure at E12.^14,22,42^ To better understand the time course of transcriptional dysregulation in frontal cortex, we performed RNA-seq of both whole frontal cortex in C57Bl/6J mice, as well as in cortical excitatory neurons from INTACT mice, at both pre- and post-natal time points. Our data from whole frontal cortex samples showed that the major transcriptional alterations caused by PIC-MIA occurred around the time of birth (P0) (Supplementary Fig. 2a, Supplementary Table 5). When evaluating our excitatory neuron-specific RNA-seq analyses in the context of other studies, we identified 28 strongly correlated overlapping DE genes at E14.5^22^ (Supplementary Fig. 2b). However, when compared to single-cell transcriptomic studies of mouse development (Supplementary Fig. 2c,d), we found *H2afz* to be dysregulated at E15 across cortical cell types (callosal projection, corticothalamic projection, and migrating subventricular zone neurons) both within and between MIA studies.^42,43^

Focusing on excitatory neurons, our cell type-specific RNA-seq data confirmed that transcriptional effects of PIC-MIA emerge by E15 and persist at P0 and P13 (see metadata in supplementary tables). Excitatory neurons from the frontal cortex had profound changes in gene expression due to PIC-MIA (Supplementary Table 6) at P0 (2113 up- and 2295 down-regulated genes, FDR<0.1) (Fig. 3a center). By contrast, far fewer genes were altered at E15, three days after the PIC injection (52 up- and 80 down-regulated genes) (Fig. 3a left). There were also relatively few significant DEGs at P13 (24 up- and 31 down-regulated genes) (Fig. 3a right).

**Fig. 3.**
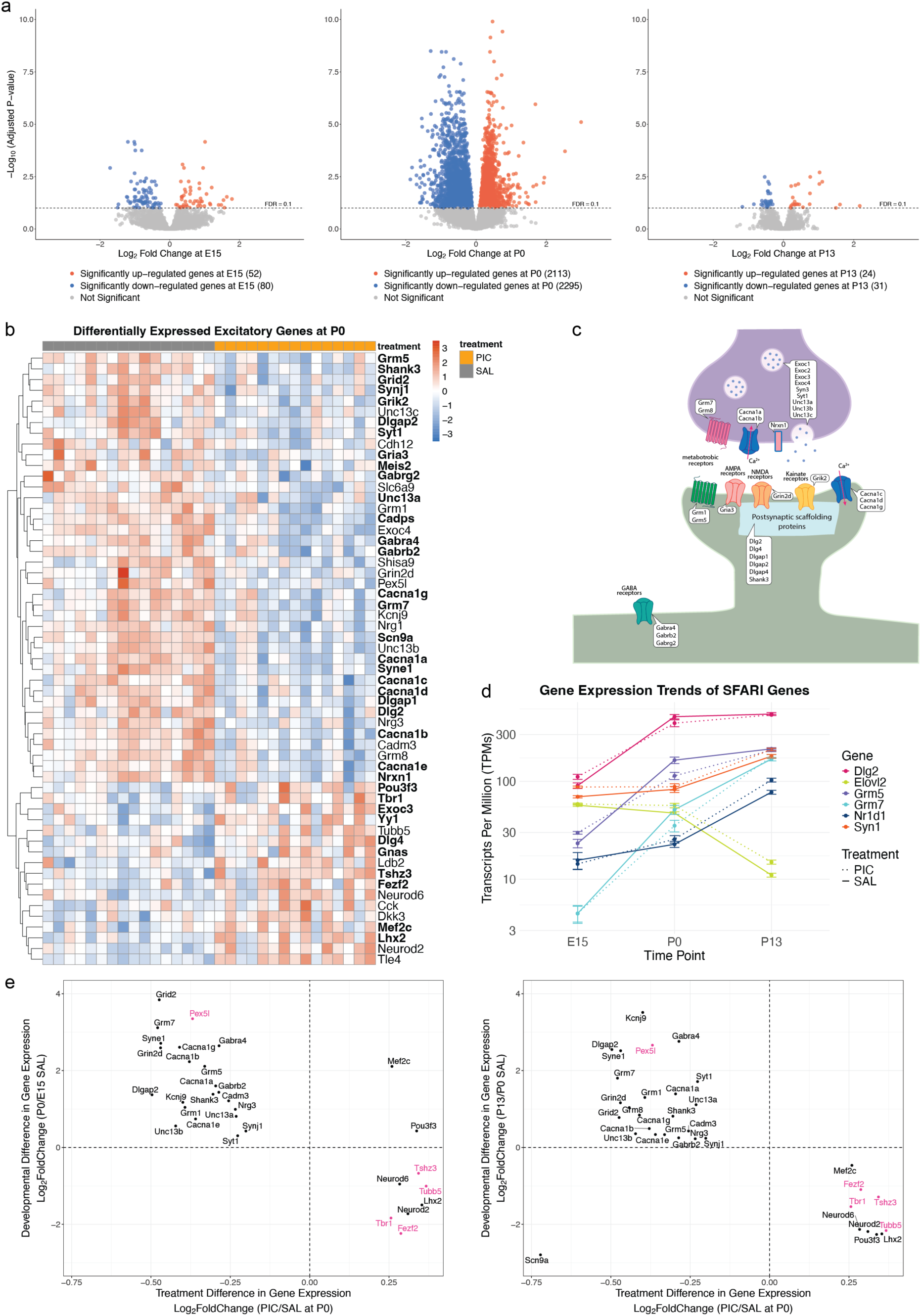
Differentially expressed (DE) genes associated with PIC-MIA surge at birth. **a**. Volcano plots with a log fold change of gene expression against adjusted p-value showing the transcriptional changes in FANS sorted excitatory neuron population in the frontal cortex of offspring of PIC vs. SAL at E15 (left), P0 (center), and P13 (right). Orange points are up-regulated and blue are down-regulated genes in PIC. Grey dots are not significantly differentially expressed genes. **b.** Distribution of excitatory neuron DE genes at P0 involved in excitatory and synaptic function. The heatmap was plotted using variance-stabilized expression values, with each gene re-scaled by Z-score normalization and ordered by hierarchical clustering. Red indicates higher expression levels while blue denotes lower expression. Genes in bold are found in the SFARI dataset. **c.** Schematic diagram of DE genes in synaptic neurons. **d.** Comparison of trends in averaged TPMs across development for selected genes from the SFARI dataset between PIC and SAL groups. Expression values were averaged within each group at each time point, and error bars indicate variability within treatment and age groups. SAL is represented by a solid line, while PIC is shown with a dotted line. Differential expression was observed for Dlg2 and Syn1 at E15, Grm 5 and 7 at P0, and Elovl2 and Nr1d1 at P13. **e.** Synaptic gene expression changes in response to MIA across developmental transitions. Scatter plots comparing treatment-induced changes in gene expression at P0 (log₂ fold change, PIC vs. SAL; x-axis) with developmental changes in gene expression from SAL group from E15 to P0 (left) and P0 to P13 (right) (log₂ fold change; y-axis). Each point represents a DE gene at P0. Genes colored in pink are enriched in deep-layer neurons.

Although the number of DEGs was modest at E15, the identity of the dysregulated genes suggested a specific impact on synaptic processes and cortical development. Up-regulated DEGs included *Grin3a, Syn3, Dlg2*, and *Unc13c,* while down-regulated genes included *Pax6, Notch1/2*, and *Sox6* (Supplementary Fig. 3a left). At P0, the down-regulation of *Scn9a* and *Npas4* could indicate altered synaptic function and excitation, while the up-regulation of *Ctsb* and *Siah1* (Supplementary Fig. 3a center) may indicate activated stress and immune responses. At P13, the up-regulation of immediate early genes (*Npas4, Arc, Fos*, *Egr1/2/4*) (Supplementary Fig. 3a right) suggests a disruption in physiological activity. Moreover, developmentally regulated genes known to play distinct roles in neuronal differentiation and function are disrupted at E15 and P0 (Supplementary Fig. 3b). These results suggest a strong effect of PIC-MIA on offspring brain development that is delayed relative to the acute immune activation, peaking around the time of birth.

PIC-MIA produced a down-regulation of Gene Ontology (Biological Process) categories associated with cell cycle regulation and an early up-regulation of genes related to synapse-related categories at E15 (Supplementary Fig. 3c) using EnrichR^37,38^ (p<0.003). Those synaptic categories were subsequently down-regulated by P0 (Supplementary Fig. 3d), suggesting that PIC-MIA induces a developmental disruption in synaptogenesis. Notably, down-regulated synaptic genes (Fig 3b,c) include synaptic adhesion molecules (*Nrxn1* and *Cadm3*), calcium channels (*Cacna1a/b/c/d/e/g*) which affect neurotransmitter release, pre- and postsynaptic metabotropic receptors (*Grm7/8* and *Grm1/5*, respectively), postsynaptic ionotropic receptors (*Gria3* and *Grik2*), as well as postsynaptic scaffolding proteins (*Dlgap1/2/4* and *Dlg2/4*). We also observed down-regulation of genes involved with GABA receptors (*Gabra4*, *Gabrb2*, *Gabrg2*) and *Slc12a5* (*Kcc2*), suggesting a delayed GABA switch in MIA offspring.

To assess the transcriptomic effects of PIC-MIA in the context of neurodevelopmental disorders, we examined the expression trends of excitatory cortical genes that have been implicated in ASD using the Simons Foundation Autism Research Initiative database.^44,45^ This database compiles autism-associated genes from genome-wide association studies, rare variant analyses in families, and functional studies. Of 1219 ASD-linked genes, we found 287 (23.54%) were differentially expressed at P0, including 60 (25.64%) with a high-confidence SFARI score of 1. Disrupted genes associated with ASD were found in all perinatal time points (Fig. 3d). For instance, *Dlg2* and *Elovl2* were DE at E15 and *Dlg2* continues to be disrupted at P0 along with *Grm5/7*. At P13, *Nr1d1* and *Syn1* were upregulated in PIC-MIA samples. Assessing the effects of MIA in genes from the SFARI dataset may serve as a critical link between disrupted developmental processes and altered excitatory cortical function.

The majority of synaptic DEGs are normally up-regulated during cortical development (Fig. 3e,f). This suggests that PIC-MIA delays normal developmental gene expression. Nevertheless, transcriptional processes are upregulated at P13 (Supplementary Fig. 3e). In addition to synaptic gene dysregulation, we found chromatin remodeling genes were disrupted at P0 (Supplementary Fig. 3f). Specifically, *Ino80c*, *Smarca5,* and *Smarcb1* are up-regulated while histone-modifiers like *Kmt2a* and *Hdac8/9* are down-regulated. Disrupted chromatin remodelers may impair synapse-related gene programming, potentially contributing to the altered trajectory of synaptogenesis in PIC-MIA offspring.

### Profound alteration of DNA methylation in PIC-MIA offspring at birth

To select the litters for MethylC-seq (whole genome bisulfite sequencing) analysis, we assessed treatment response by gene expression profiles. We used principal component scores and linear discriminant modeling to identify the pups with the strongest PIC vs. SAL classification scores (see sample selection in methods). Excitatory neuron-specific MethylC-seq analyses were performed for excitatory neurons from single pups (n=6 mice from 3 litters/group for P0 and P13, see metadata). We identified PIC-MIA differentially methylated regions (miaDMRs) with a significant difference in mCG between PIC-MIA offspring and SAL-treated controls (>10% difference in mCG). We found 2,607 miaDMRs in excitatory neurons at P0, long after the acute phase of PIC treatment, while fewer methylation differences were observed at P13 (196 miaDMRs) (Fig. 4a). The mCG changes at P0 included 1,548 regions with lower mCG and 1,069 with higher mCG in PIC-MIA compared to controls (Fig. 4a and Supplementary Table 7).

**Fig. 4.**
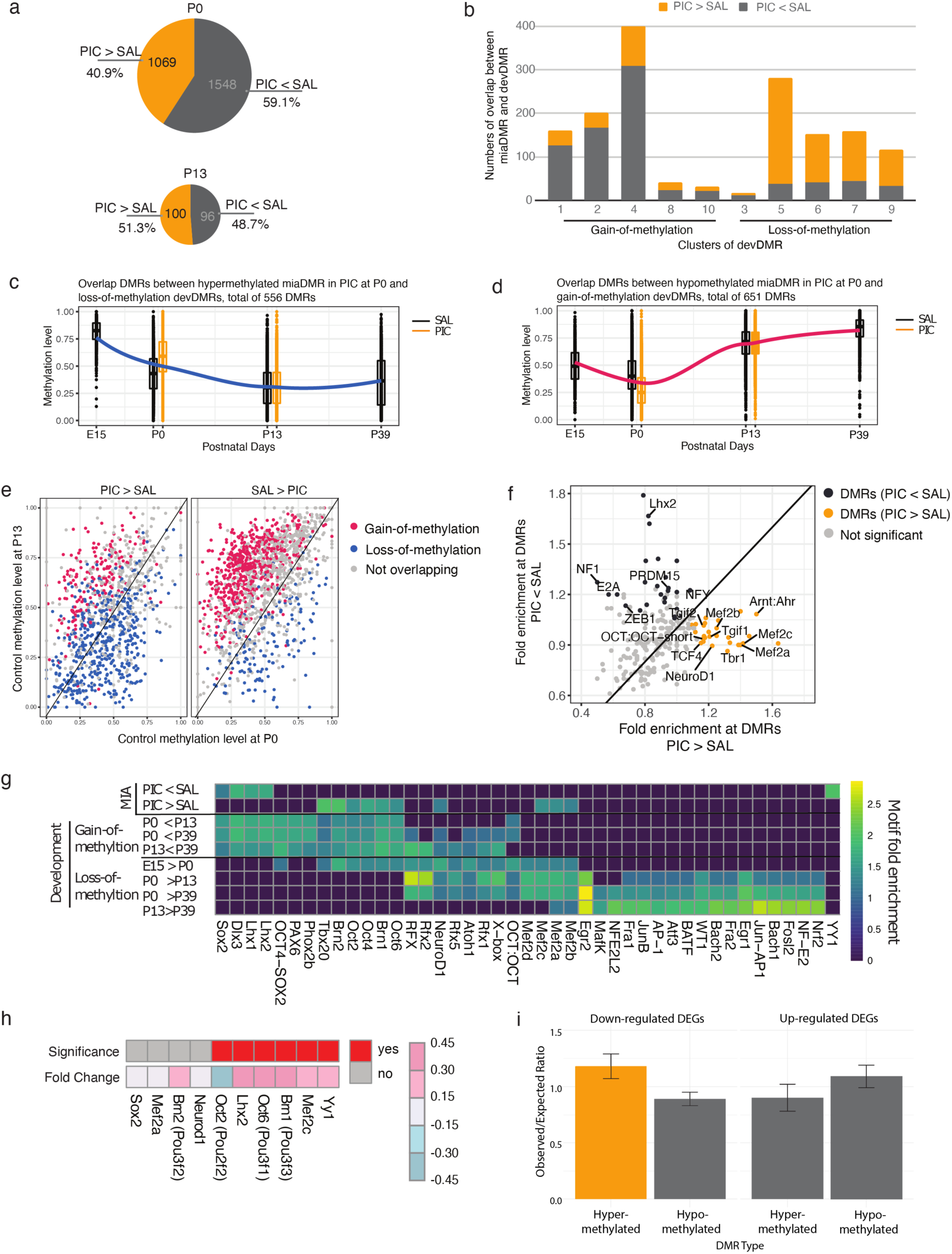
Differential DNA methylation that was altered by PIC-MIA was profound at birth. **a.** Numbers of differentially methylated regions between offspring of PIC and SAL injection mice (miaDMRs) at P0 and P13. PIC > SAL, hypermethylated in PIC. PIC < SAL, hypomethylated in PIC. **b.** Enrichment of miaDMR among devDMR clusters. The clusters of developmental loss-of-methylation are enriched for miaDMRs that are hyper-methylated in PIC. While the clusters of developmental gain-of-methylation are enriched for miaDMRs that are hypo-methylated in PIC. **c.** Dynamics of methylation levels of hyper-methylated miaDMRs (PIC > SAL) or **d.** hypo-methylated miaDMRs (PIC < SAL) that overlap with devDMRs. From the perspective of normal development, the methylation levels of miaDMRs was delayed at birth but returned to normal in postnatal day 13. **e.** The joint distribution of control methylation level of P0 and P13 in miaDMRs. The miaDMRs that overlap with devDMRs are marked with blue or red color, and miaDMRs that do not overlap with devDMRs are gray dots. For hyper-methylated miaDMRs (PIC > SAL), the distribution shifts towards P0, indicating enriched loss-of-methylation (blue dots) during development. Meanwhile, the distribution shifts towards P13 in hypo-methylated miaDMRs (PIC > SAL), meaning that they are enriched in gain-of-methylation (red dots) during development. **f.** Enriched transcription factor binding motifs in miaDMRs. Black/orange dots are significantly enriched transcription factor binding motifs in hypo-/hyper-methylated miaDMRs. **g.** The TF binding motifs in miaDMRs are also enriched in devDMRs. The color indicates fold enrichment level which normalized the percentage of target enrichment to the background signal, the higher value indicates more significantly enriched. Lhx1/2, Dlx3, and Sox2 were enriched in both hypo-methylated miaDMRs and gain-of-methylation devDMRs. POU domain families (Brn1-2 and Oct2,4,6) were enriched in both hyper-methylated miaDMRs and gain-of-methylation devDMRs. Mef2a-c were enriched in both hyper-methylated miaDMRs and loss-of-methylation devDMRs. **h.** The gene expression fold-change of TF binding motifs. Lhx2, Yy1, Mef2c, and two of POU domain families (Brn1 and Oct6) were up-regulated in PIC samples. Lhx2 is hypo-methylated and up-regulated in gene expression of PIC at P0. It is developmental hyper-methylated (P13 > P0) and developmental down-regulated in gene expression. **i.** Enrichment analysis between miaDMRs and miaDEGs. Bar plots show the hyper- or hypo-methylated DMRs on the x-axis and the observed/expected ratios of overlaps between miaDMRs and miaDEGs on the y-axis. Each bar represents the enrichment of up- or down-regulated DEGs in hyper- or hypo-methylated DMRs. Yellow bars indicate significantly enriched pairs of miaDEGs and miaDMRs.

Next, we compared miaDMRs with differentially methylated regions during development (devDMRs). Among 729 miaDMRs identified at P0 that overlapped with loss-of-methylation devDMRs, the majority (556, 76.3%) were hyper-methylated in PIC-MIA (Supplementary Table 8). The miaDMRs had a substantial correlation with specific devDMR clusters (Fig. 1b), with regions that lose methylation during normal development (devDMR clusters 3, 5, 6, 7 and 9) mainly overlapping miaDMRs that gain methylation (PIC > SAL) (29%, 86%, 71%, 73%, and 71%, respectively; Fig. 4b right panels and Supplementary Table 8). On the other hand, for the 838 miaDMRs that overlapped with gain-of-methylation devDMRs, 651 (77.7%) were hypo-methylated DMRs in PIC-MIA. Regions that gain methylation during development (devDMR clusters 1, 2, 4, 8, and 10) overlapped miaDMRs that lose methylation (78%, 83%, 78%, 56%, and 69%, respectively; Fig. 4b left panels and Supplementary Table 9).

To visualize the dynamics of hypermethylated (PIC > SAL) or hypomethylated (PIC < SAL) miaDMRs during normal development, we plotted the 556 overlapping loss-of-methylation devDMRs and hypermethylated in PIC-MIA at P0 throughout the development time points. PIC-MIA delayed loss-of methylation of devDMRs in excitatory neurons at birth (Fig. 4c). A similar pattern was observed for the 651 overlapping gain-of-methylation devDMRs and hypomethylated in PIC-MIA at P0 (Fig. 4d). To further illustrate how these miaDMRs align with normal postnatal methylation trajectories, we plotted the control (SAL) methylation levels at P0 and P13 for each DMR (Fig. 4e). Most hypermethylated miaDMRs (PIC > SAL) correspond to regions that undergo developmental loss of methylation from P0 to P13 (blue), indicating that the delay of normal hypomethylation persists at P0. Conversely, hypomethylated miaDMRs (PIC < SAL) tend to occur in regions that normally gain methylation (red), suggesting a disruption of these maturation-associated increases (Fig. 4e). However, these effects disappeared when animals reached the second postnatal week (Fig. 4c,d), suggesting that PIC-MIA affected the DNA methylation program during the early postnatal period before the activation of Dnmt3a during the second postnatal week.^17^

To interpret the potential regulatory role of miaDMRs at P0, we performed transcription factor binding motif (TFBM) enrichment analysis and compared the results with enriched TFBM for devDMRs (Supplementary Table 3). Results of these analyses further suggest that PIC-MIA produces a delay in the normal maturational program of excitatory neurons (Fig. 4f). Binding sites for TFs involved in neuronal maturation (e.g. *Mef2* and *Neurod* family factors) appear hypermethylated and thus repressed in MIA. In contrast, TFBMs that should be silenced at birth (such as *Sox*, *POU*, and *Lhx* families) remained demethylated (Fig. 4g, Supplementary Table 10). The expression level of TFs involved in early neurodevelopment (*Lhx2*, *Brn1*, and *Yy1*) and neuronal maturation (*Mef2c*) were significantly dysregulated in PIC-MIA (Fig 4h). *Lhx2* and *Yy1* expression was increased and their binding sites demethylated, suggesting an increased function of these TFs in PIC-MIA offspring. On the other hand, Oct2 expression was downregulated and its binding sites hypermethylated, consistent with diminished activity of this TF in PIC-MIA offspring (Fig. 4g,h).

To further understand the regulatory dynamics between the miaDMRs and DEGs, we performed an enrichment analysis to identify the DE genes at P0 that were associated with DMRs at P0. Our results showed that hyper-methylated DMRs were significantly associated with down-regulated genes (observed/expected ratio 1.18, p=0.03) (Fig. 4i, Supplementary Table 11); we found no significant association between up-regulated genes and hypo-methylated DMRs (p=0.232).

### Cell type-specific roles of miaDMRs

To further refine our interpretation of the cell type-specific role of regulatory elements whose methylation is affected by PIC-MIA, we compared miaDMRs with single-cell resolution DNA methylome profiles from the mouse and human frontal cortex and mouse motor cortex^29,46^ (Fig. 5). Hyper-methylated miaDMRs (Supplementary Table 12) were significantly enriched in cell type-specific DMRs with low methylation in mouse frontal cortex deep-layer excitatory neurons^29^ (cluster DL2; Fig. 5a), but not in mouse motor cortex. Hypo-methylated miaDMRs were enriched in regions that were hypomethylated specifically in GABAergic neurons, including CGE-derived Lamp5 and Vip neurons in motor cortex (Fig. 5b); in Ndnf-1, Ndnf-2, and Vip cell types in mouse frontal cortex (Fig. 5c); and Ndnf, Nos, Vip-1, and Vip-2 cell-types in human frontal cortex (Fig. 5d). This pattern suggests that regulatory regions used by GABAergic neurons, normally methylated and repressed in excitatory neurons, remain anomalously hypomethylated in excitatory neurons of PIC-MIA mice. To further understand whether miaDMRs could affect the gene expression of specific cell types, we performed enrichment analyses between the cell-type-specific miaDMRs and the miaDEGs. We found a significant association between layer 5 and 6 neuron DMRs and hypo-miaDMRs that overlapped with up-regulated DE genes (Fig. 5e and Supplementary Table 13).

**Fig. 5.**
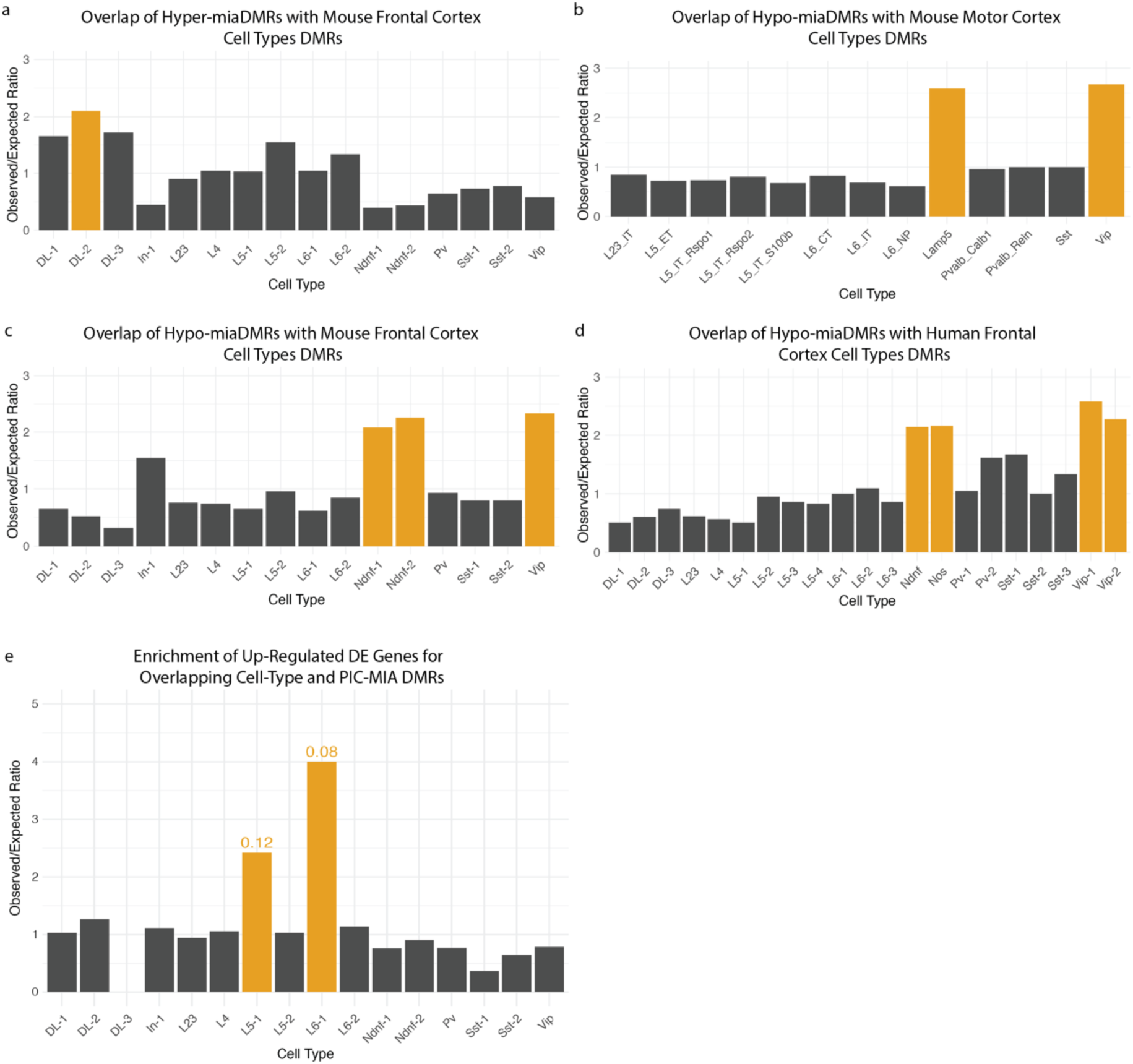
Enrichment analyses between cell-type DMRs, miaDMRs, and DE genes. Bar plots display enrichment results, with cell-type DMRs on the x-axis and observed/expected ratios of overlaps on the y-axis. Cortical cell-type DMRs were obtained from Luo et al.^29^ and Yao et al.^46^ (**a–d**) Enrichment of hypo- and hyper-miaDMRs in cell type DMRs. Each bar represents the enrichment of hypermethylated (hyper) or hypomethylated (hypo) miaDMRs in DMRs associated with specific cell types. Cell types highlighted by yellow bars have observed/expected ratios>2 and adjusted p-values<0.01. **e.** Analyses of observed/expected ratios of up-regulated differentially expressed genes for overlapping cell-type and PIC-MIA DMRs. Cell types highlighted by yellow bars have observed/expected ratios>2 and adjusted p-values<0.15. Significant enrichment was only observed in up-regulated DE genes associated with hypo-miaDMRs.

### Layer-specific effects of maternal immune activation on neuronal physiology

The DNA methylation and transcriptomic results described above suggested differential effects on cortical layer-specific neurons. To study such effects, we used whole-cell patch-clamp recordings to measure electrophysiological parameters indicative of neuronal maturation. These recordings were conducted during critical postnatal stages (P6-9 and P14-17) in layer 5 (L5; Fig 6, Supplementary Fig. 5a,d) and layers 2/3 (L2/3; Supplementary Fig. 5e,f) prelimbic pyramidal neurons. Our findings shed light on the differences in neuronal excitability and synaptic activity in the brains of offspring from dams treated with either saline or PIC.

**Fig. 6.**
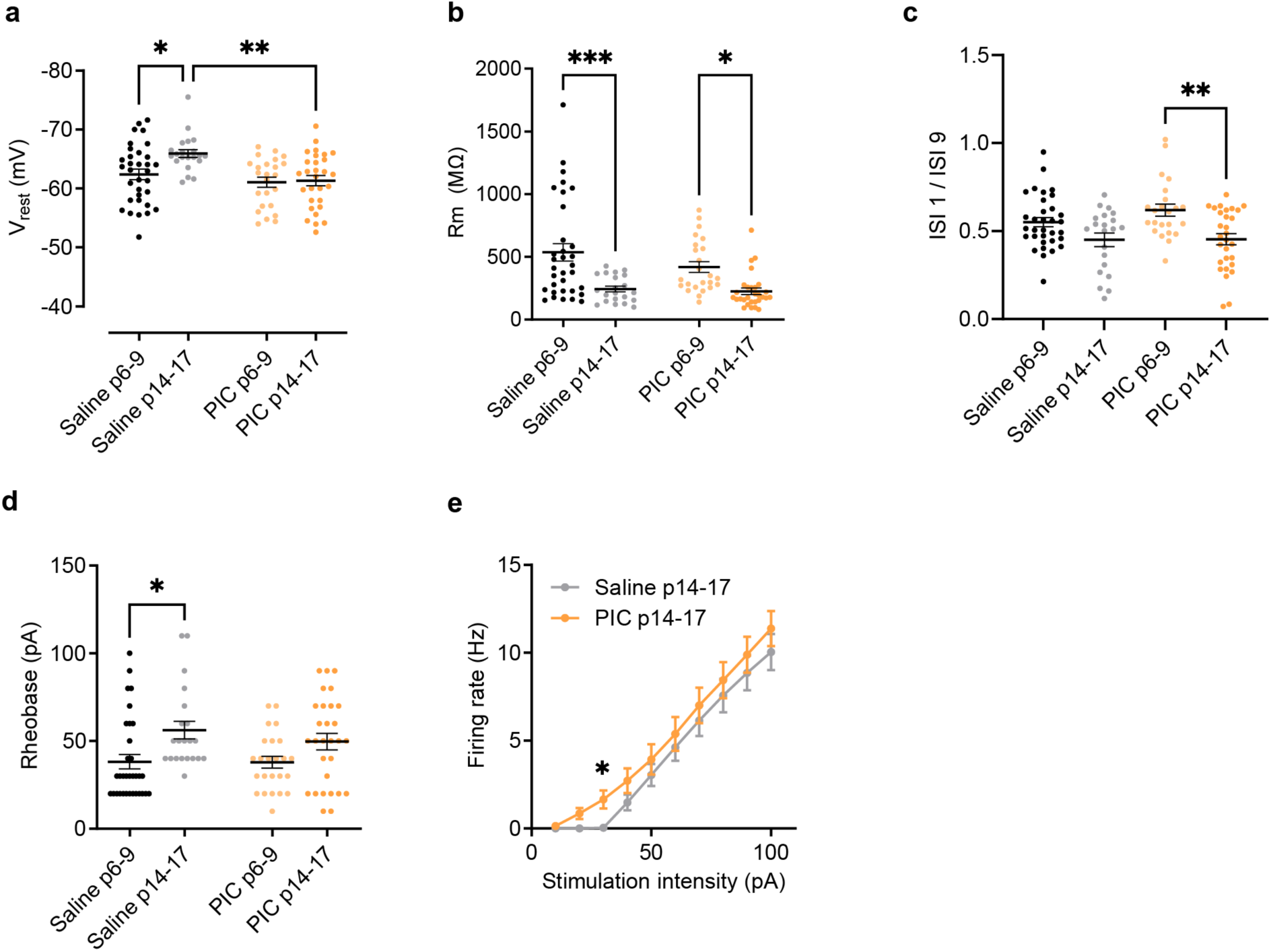
Effect of maternal immune activation on the postnatal maturation of L5 pyramidal neurons. Comparison of saline- and PIC-treated groups at two developmental time points (P6-9 and P14-17) across passive (a, b) and active intrinsic properties (c-e). **a.** Resting membrane potential (Vrest) exhibited developmental hyperpolarization in saline-treated mice but remained unchanged in PIC-treated mice. **b.** Membrane resistance (Rm) decreased over time in both conditions. **c.** Adaptation, measured as the ratio of the first over the ninth interspike interval (ISI1/ISI9), increased with age (i.e., the ratio decreased, indicating stronger adaptation), reaching statistical significance only in PIC-treated mice. **d.** Rheobase increased during development, with a significant effect observed in saline-treated mice but not in PIC-treated mice. **e.** Input-output curves revealed a selective difference near threshold (30 pA) between saline- and PIC-treated mice at P14–17. Data points represent individual neurons, with mean values indicated by horizontal bars. Error bars denote SEM. Statistical significance: *p<0.05, **p<0.01, ***p<0.001, ****p<0.0001.

Our findings highlighted a temporal evolution in several electrophysiological parameters of L5 pyramidal neurons within both groups. Alterations in maturation markers, such as the advancement to more hyperpolarized resting membrane potentials (RPM), were significantly reduced, in the PIC-MIA group. Specifically, the RMP of L5 neurons shifted significantly (p<0.05) between time points from −62.4 ± 0.9 mV (n = 33 cells) to −65.9 ± 0.7 mV (n=21 cells) in saline mice, but it remained stable in PIC-MIA mice (−61.1 ± 0.9, n = 24 cells, and −61.4 ± 0.9, n = 29 cells) (Fig. 6a). This shift was further reflected as a significant difference (p<0.05) between treatment conditions at the later time point. Membrane resistance decreased over time in both control and PIC-MIA mice (Fig. 6b), accompanied by a shift in action potential threshold to more negative values (Supplementary Fig. 4a), a reduction in action potential half-width (Supplementary Fig. 4b), and an increase in action potential amplitude (Supplementary Fig. 4c), without significant differences between treatment conditions. However, specific parameters, such as adaptation (ISI 1/ISI 9) (Fig. 6c) and rheobase (Fig. 6d) evolved differently between groups. This included a developmental shift in rheobase that was statistically significant (p<0.05) only in control conditions (from 38.2 ± 4.1 pA, n = 33 cells to 56.2 ± 5.0 pA, n = 21 cells vs PIC-MIA: 37.9 ± 3.4, n = 24 to 49.7 ± 4.7 pA, n=29) (Fig. 6d). Despite the lack of a significant difference in rheobase between saline and PIC-MIA treatments when analyzed in isolation, input-output curves revealed a selective difference near threshold at the later developmental time point. Specifically, repeated-measures ANOVA showed that while firing rates were largely similar between conditions, a significant difference existed at the 30 pA intensity level (Fig. 6e). This suggests that although the absolute rheobase values remained comparable, subtle differences in neuronal excitability or spike initiation dynamics may have altered the transition from subthreshold to suprathreshold firing. Note that, although subtle, the alterations induced by maternal immune activation upon the typical postnatal maturation trajectory of L5 pyramidal neurons occurred in aspects related to excitability. Altogether, these differences hint at a detrimental effect of maternal immune activation on neuronal excitability during early postnatal development, potentially compromising their signal specificity and efficiency.

We also recorded miniature excitatory postsynaptic currents (mEPSCs) to assess differences in synaptic function and postsynaptic receptor properties in both L5 and L2/3 neurons. In L5 neurons, both control and PIC-MIA groups exhibited a maturation-dependent increase in mEPSC frequency (Fig. 7a). Interestingly, while mEPSC amplitude did not change significantly across time points under control conditions, L5 neurons in the PIC-MIA group showed a time-dependent decrease in amplitude (Fig. 7b). This reduction may be linked to the finding that, in the youngest cohort (P8–P9), mEPSCs recorded from PIC-MIA L5 neurons were significantly larger (p<0.05) than those in the control group (Fig. 7b), suggesting a maturational delay in synaptic properties. The average mEPSC amplitude at this time point was 13.0 ± 0.5 pA in PIC-MIA neurons (n= 14 cells), compared to 11.2 ± 0.5 pA (n = 9 cells) in control neurons. Analysis of mEPSCs in L2/3 neurons revealed that the frequency increased, and amplitude decreased between developmental time points in both control and PIC-MIA neurons (Fig. 7c,d).

**Fig. 7.**
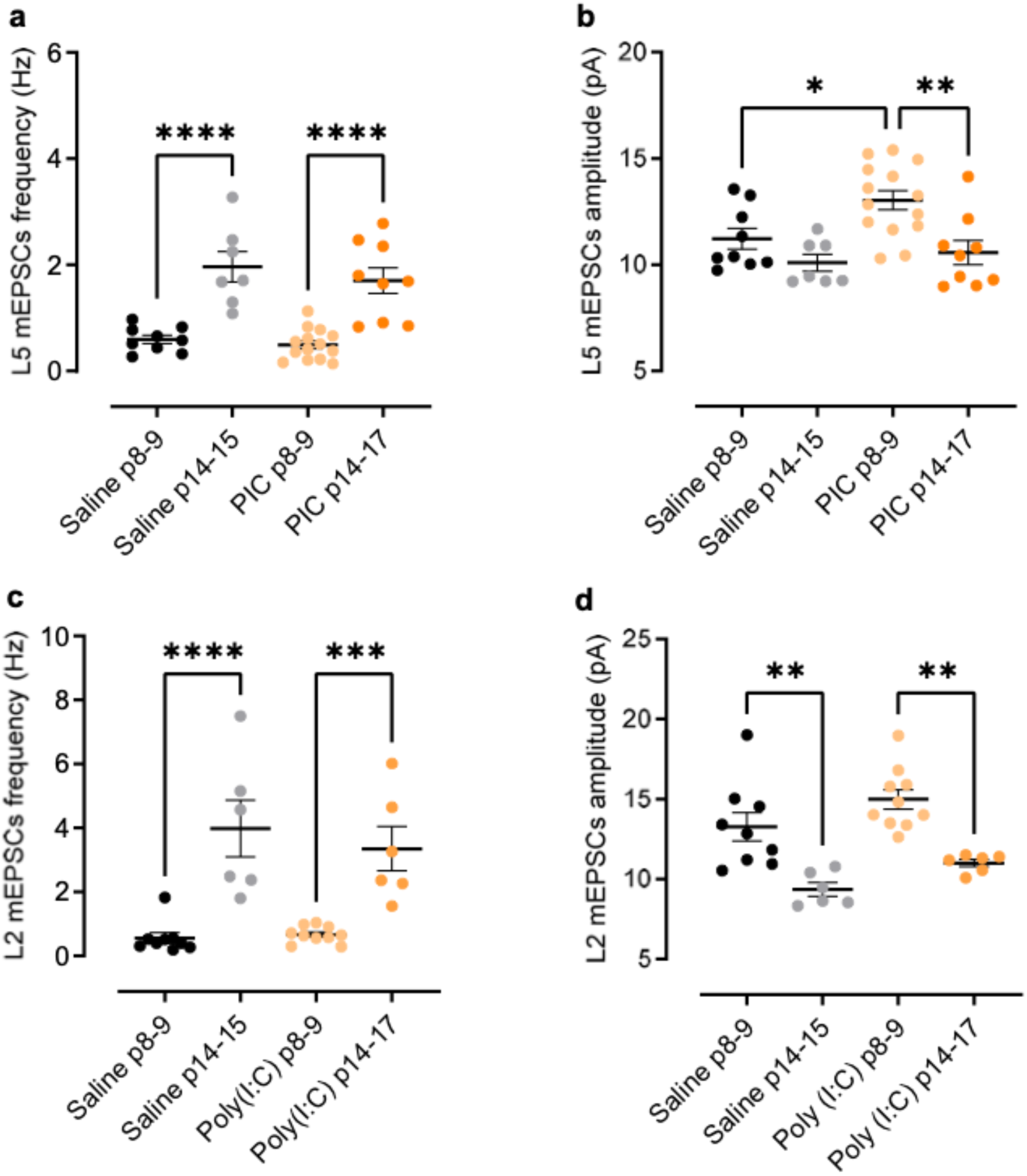
Effect of maternal immune activation on the postnatal maturation of excitatory synaptic inputs. mEPSCs were recorded from L5 (a,b) and L2/3 (c,d) pyramidal neurons at two developmental time points (P8-9 and P14-17). mEPSCs frequency increased with age in both L5 (**a**) and L2/3 (**c**) neurons of saline- and PIC-treated mice. mEPSCs amplitude recorded in L5 (**b**), but not L2/3 (**d**) neurons, was significantly higher at P8-9 in PIC-treated mice compared to saline-treated mice. No age-dependent change in mEPSCs amplitude was observed in saline-treated mice, while a significant decrease was observed between time points in L5 neurons of PIC-treated mice. Note that such a reduction suggests a potential developmental delay in synaptic maturation. Data points represent individual neurons, with mean values indicated by horizontal bars. Error bars represent SEM. Statistical significance: *p<0.05, **p<0.01, ***p<0.001, ****p<0.0001.

Extending our analyses to characterize the electrophysiological properties of L2/3 pyramidal neurons allowed us to assess whether the observed developmental impacts of PIC-MIA were layer-specific or exhibited a broader effect across cortical layers. In both L5 and L2/3 neurons, we found a significant maturation of electrophysiological parameters within both groups. Contrasting with L5 neurons, however, PIC-MIA did not affect the maturational shift in L2/3 neurons’ RMP or rheobase, which was similar in both groups (Supplementary Fig. 4e,f). At P7-9, the RMP in treated L2 neurons was −60.3 mV ± 1.2 mV (n = 31 cells), compared to −60.9 ± 1.3 mV (n = 23 cells) in control conditions. By P14-17, the RMP had become more hyperpolarized in both conditions (control: −66.7 ± 1.0 mV, n = 32 cells; PIC-MIA: −67.8 mV ± 1.3 mV, n = 24 cells). Taken together, these results suggest that the impact of maternal immune activation is greater in deep layers of the cortical column.

## Discussion

Our findings uniquely integrate multiple layers of biological data—DNA methylation, transcriptional profiles, and electrophysiological outcomes—to provide a comprehensive understanding of cortical neuron maturation disrupted by maternal immune activation. We observed coordinated disruptions wherein altered DNA methylation at key developmental regulatory regions was closely aligned with transcriptional dysregulation of genes essential for neuronal maturation, synaptogenesis, and circuit formation. Corresponding electrophysiological analyses demonstrated functional consequences of these epigenetic and transcriptional disruptions, particularly in deep-layer cortical neurons. As previously shown for whole tissue^17,47^ and single-cells,^48^ excitatory neuron maturation in the frontal cortex entails a profound epigenomic rearrangement starting at mid-gestation and lasting through the first two postnatal weeks. Normal developmental methylation patterns during this period are defined by a profound alteration in the methylation of putative enhancer regions (as defined by gain or loss of H3K27ac peaks) before and after birth, including transcription factor binding sites of neurodevelopmental regulators (*Neurog2*, *Lhx2*). Developmental methylation changes aligned with neural progenitor downregulation and proneural gene upregulation at birth, consistent with prior studies.^49^ In this context, we observed that normal developmental patterns were dramatically altered by PIC-MIA.

Epigenomic disruption of the offspring following maternal immune manipulation was most pronounced at birth, long after the maternal immune activation occurred, yet the transcriptomic differences observed at E15 already point to early developmental effects. Similar to single-cell studies,^42,43^ our data suggest an initial push towards cell differentiation, followed by increased protein synthesis at P0. The down-regulation of the cell cycle (*Ccnd1*, *Cdk1/6*) and neurogenic (*Neurog2, Notch1/2*, *Pax6*, *Sox6*) regulators in our E15 data suggest halted neural progenitor proliferation and premature differentiation. Despite this early shift, epigenomic signatures at P0 suggest developmental delay with stalled synaptic development marked by deficits in scaffolding proteins (*Dlg2/4*, *Shank3*) and glutamatergic transmission (*Grm1/5/7/8*, *Gria3*, Grin2d). Consistently, DNA methylation patterns also reflected disrupted developmental timing. Transcription factor binding sites such as *Sox2*, *Dlx3*, and *Lhx1/2*, which normally gain methylation during development, remained hypomethylated in miaDMRs at P0. Binding sites for factors that normally lose methylation with development, such as *Mef2a-c*, remained hypermethylated in miaDMRs. Collectively, this pattern suggests a coordinated disruption in both transcriptional and epigenomic regulation following PIC-MIA. Although our results indicate a strong correlation between altered DNA methylation and transcriptional dysregulation, establishing direct causality was beyond our current scope. Future experiments utilizing targeted epigenome editing or CRISPR-based manipulation of identified methylation sites could confirm the direct regulatory roles of specific methylation changes in driving observed transcriptional alterations.

While DNA methylation alone does not dictate gene expression, its interplay with chromatin remodelers can shape transcriptional landscapes. We found significant disruptions in genes involved in chromatin regulation such as *Smarca5* and *Smarcb1*, alongside histone-modifying enzymes like *Kmt2a/b* and *Hdac8*,^50^ which exhibited altered expression at P0. Additionally, *Yy1* has been proposed as a structural regulator of enhancer-promoter loop^51^ where the loss of methylation of its binding sites—as we found in PIC-MIA—may promote changes in enhancer-promoter interactions. As in single-cell studies,^42,43^ we found that histone variant *H2afz* was dysregulated following PIC-MIA, suggesting that impaired nucleosome stability and enhancer accessibility may contribute to broad epigenomic dysregulation of transcriptional control.^52^ Given that chromatin remodelers play critical roles in the transcriptional regulation of neuronal development,^53^ disrupted expression may underpin gene dysregulation observed in PIC-MIA offspring.

Although PIC-MIA effects on DNA methylation and RNA transcription were observed mostly after birth, these epigenomic changes–at the time when cortical circuits are formed– could lead to the miswiring of the developing circuitry and behavioral alterations later in life. Indeed, our findings demonstrate subtle, yet significant alterations in excitability and synaptic properties in L5, but not in L2 neurons, emphasizing the layer-specific impact of PIC-MIA. The heightened excitability of L5 neurons could be linked to the up-regulation of activity-dependent immediate early genes (*Arc*, *Egr1/2/4*, *Fos*, and *Npas4*) observed at P13, potentially compensating for disrupted deep layer network activity. Beyond immediate early gene activation, analysis of miaDMRs and cell-type-specific DMRs further supports an enrichment of epigenomic changes in deep-layer neurons. Genes predominantly expressed in deep layers (*Ntsr1, Cck*, *Fezf2*, *Ldb2*, *Tle4*, *Tbr1*, and *Pex5l*)^48^ were dysregulated at P0 which may point to altered circuitry formation. Deep layer cortical neurons play a fundamental role in the establishment of intra and extra-cortical circuitries during the first postnatal weeks in mice, and alterations in their maturation are expected to disrupt the program of circuit formation.^54,55^ In this line, our electrophysiological analyses indicated that layer 5 (L5) pyramidal neurons were particularly susceptible to maternal immune activation, whereas L2/3 neurons showed less pronounced alterations. This layer-specific vulnerability might reflect the distinct developmental timelines and connectivity patterns inherent to deep-layer neurons, highlighting a critical window in deep-layer cortical neuron maturation and circuit formation disrupted by prenatal immune challenges. Our present findings highlight the complexity of cortical development and the importance of a detailed understanding of how prenatal immune challenges affect neuronal maturation across different cortical layers.

Assessing DE genes against the SFARI dataset revealed strong links between ASD-associated genes and PIC-MIA-induced transcriptional changes in excitatory neurons. We concentrated on the male offspring given the high male/female ratio for ASD in human studies.^56^ PIC-MIA disrupted genes that were both associated with ASD and developmentally regulated, revealing insights into mechanisms driving altered brain maturation. We found disruptions in synaptic genes (*Gria3*, *Grin2d*, *Shank3*, *Grm1/5/7/8*) critical for maintaining the excitation/inhibition balance in neural circuits that have been implicated in ASD.^57^ *Grm5* expression is decreased in ASD individuals,^58^ and clinical and functional data demonstrated that both autosomal dominant and recessive inheritance of GRM7 mutation can cause disease spectrum phenotypes through ASD.^59,60^ Furthermore, genes with dysregulation at birth persisting into adulthood (*Csmd3*, *Grm7, Kcnk1*, *Polr2a*, and *Ssr2*),^8^ may reflect long-term consequences of PIC-MIA. Additionally, the *Egr* family which is dysregulated at P13 has been implicated in atypical synaptic expression in ASD.^61^ Collectively, these findings suggest disrupted processes during circuit formation that may contribute to a predisposition to ASD.

Although PIC-MIA studies have demonstrated neurodevelopmental alterations resembling ASD phenotypes, it is critical to emphasize that these effects are mediated by elevated levels of pro-inflammatory cytokines such as IL-6.^4^ The cytokine levels post-PIC injection are comparable to those produced during a cytokine storm due to influenza or other serious infections.^62^ Such levels of proinflammatory cytokines are not produced during common cold infections or after immunizations. In the context of immunizations, the innate immune response induces a modest cytokine release, which subsequently activates a cascade of events that drive antibody production to neutralize the pathogen.^63^ While the cytokine release following immunization may raise concerns about the relationship between immunological exposure and ASD, prenatal maternal influenza immunization has not been associated with ASD risk.^64,65^

In summary, PIC-MIA had profound and lasting effects on the development of excitatory cortical neurons. Atypical development during the transition between embryonic and adult programming impacts the formation of cortical circuits necessary for proper function.^54,55,66^ Additional studies are necessary to elucidate the mechanisms underlying electrophysiological alterations following PIC-MIA and the functional consequences for brain development and behavior. Further investigation into the effects of PIC-MIA on other cell types in the brain may reveal how disrupted coordination between neuronal and non-neuronal cells contributes to increased neurodevelopmental disorder susceptibility. Furthermore, assessing changes induced by PIC-MIA at more fine-grain time points may pinpoint the temporal effects on developmental processes. Integrated multi-omic approaches and circuit-level analyses will be essential to elucidate the mechanisms by which PIC-MIA alters neurodevelopment and provide a framework to inform potential therapeutic targets and interventions.

## Limitations and future directions

This study presents an assessment of the effects of PIC-MIA on DNA methylation and transcriptional landscapes in developing mouse frontal cortex excitatory neurons. Due to intrinsic variability associated with maternal immune activation, we employed stringent selection criteria based on maternal weight change and principal component analysis of offspring gene expression to identify representative samples. While necessary to minimize variability, this approach introduces potential selection bias, which we acknowledge may limit generalizability. Future studies using larger cohorts could address this limitation. Further, the current study exclusively analyzed male offspring due to the higher incidence of autism spectrum disorders in males. We recognize that maternal immune activation could have distinct effects in female offspring^22^ and suggest future studies that explicitly evaluate sex differences to fully characterize the vulnerability and resilience to prenatal immune challenges. Although our study identifies significant molecular and electrophysiological alterations in cortical excitatory neurons following maternal immune activation, direct behavioral assessments were beyond the scope of this investigation. Establishing direct links between these observed cellular and molecular changes and specific behavioral phenotypes relevant to autism or schizophrenia is essential for understanding clinical implications. Future studies incorporating behavioral analyses, such as social interaction, cognitive flexibility, and anxiety-related behaviors, will be necessary to determine how these molecular and physiological disruptions translate into functional outcomes

## Materials and Methods

### Maternal Immune Activation model for disruption of neurodevelopment

All animal procedures were conducted following the guidelines of the American Association for the Accreditation of Laboratory Animal Care and were approved by The Salk Institute for Biological Studies and the University of California San Diego Institutional Animal Care and Use Committees. Animals were maintained under a 12-h light/12-h dark cycle in a temperature-controlled room with *ad libitum* access to water and food until euthanasia. The temperature in the animal facility was maintained within the range of 20 to 22 °C, while the humidity levels varied between 35 and 60%.

To be able to isolate pyramidal neuron nuclei for DNA methylation and transcription studies, we used a mouse line that carries a Cre-dependent nuclear envelope protein fused to GFP (INTACT mouse line, B6.129-Gt(ROSA)26Sor^tm5(CAG-Sun1/sfGFP)Nat^/MmbeJ,^23^ backcrossed to C57Bl/6J for 9 generations. ClSun1-GFP^+/+^) and crossed it to a cell-type specific Cre-line (Neurod6^tm1(cre)Kan^ = Nex-Cre). This will produce nuclei expressing GFP in their nuclear envelope that can be isolated by fluorescence-activated nuclei sorting (FANS).

For the Poly(I:C) induced MIA model (PIC-MIA), male INTACT or C57BL/6J mice were single-housed for 3 consecutive days. On the fourth day, two 10-week-old female mice were introduced to the age-matched males and left undisturbed for three days. After three days females were single housed for the remainder of pregnancy. Dams were checked daily for the presence of a seminal plug, which when found was recorded as embryonic day (E) 0.5. The weight of dams was recorded every day for pregnancy development. On E12.5 female mice received an intraperitoneal (IP) injection of 20 mg/kg Poly(I:C) acid potassium salt (P9582, Lot number 095M4085V, Sigma-Aldrich, St. Louis, MO) in 0.9% saline. This lot of PolyI:C administered at a dose of 20 mg/kg (IP) to naive female mice increased plasma IL-6 levels (Saline < 15.6 pg/ml; Poly(I:C)= 3749.6 ± 1326.2 pg/mg protein; n=3/group), confirming effectiveness. The 20 mg/kg dose of Poly(I:C) administered on E12.5 was previously shown to induce a maternal inflammatory response and produce altered behaviors in offspring.^8,67,68^ Control mice received 0.9% saline injections. The frontal cortex of the PIC- or SAL-exposed C57Bl/6J male mice were harvested at embryonic day 14.5 (E15), postnatal day 0 (P0) and 10 weeks (P70) for RNA sequencing (Fig. S3) or at P8-P17 for slice physiology (Fig. 6 and 7). For isolation of excitatory neuron nuclei from INTACT mice for RNA sequencing and whole genome bisulfite sequencing, the frontal cortex of PIC- or SAL-exposed animals were isolated at embryonic day 14.5 (E15), and postnatal days 0 (P0), 13 (P13) and 39 (P39) (Fig. 1a).

### IL-6 detection

To test for the effectiveness of Poly(I:C) batches (P9582, Sigma-Aldrich, St. Louis, MO), virgin females were injected intraperitoneally with 5 mg/kg Poly(I:C) in saline, and blood was collected 2 hours post-injection. IL-6 levels were determined in plasma following established protocols.^69^ Poly(I:C) increased plasma IL-6 levels (Saline < 15.6 pg/ml; Poly(I:C) batch 1 = 4,944 +/− 1960 and batch 2 = 6,228.2 +/− 2040 pg/ml n=3/group), confirming immunogenicity.

### Frontal cortex dissection, Nuclei isolation and flow cytometry

Frontal cortex tissue was harvested as described.^17,29^ We dissected the frontal cortex from male exposed mice at embryonic day 14.5 (E15), postnatal day 0 (P0) and 10 weeks (P70) for wild-type mice, and postnatal day 0 (P0), 13 (P13), and P39 for INTACT mice (Fig. 1a). The nuclei of excitatory neurons were isolated and collected using fluorescence-activated nuclei sorting (FANS) as described^17,29^ with the following modification: prior to FANS, nuclei were labeled with anti-NeuN-AlexaFluor647 and anti-GFP-AlexaFluor488. Nuclei were sorted as described.^33,70^ Double positive nuclei were retained for RNA-seq and MethylC-seq library preparation and sequencing (see below).

### RNA isolation and RNA sequencing

RNA was extracted by using the RNeasy Mini kit with DNAse I treatment (Qiagen, 74104) for whole frontal cortex, or by using Single-Cell RNA Purification Kit (Norgen, 51800) for nuclei. The RNA quality, RNA integrity number (RIN), was measured using the Agilent 2200 TapeStation system (Agilent Technologies, G2965AA). All samples had RIN > 7.

For whole tissue RNA, strand-specific cDNA libraries were constructed by selecting poly-adenylated mRNA transcripts using Truseq Stranded mRNA High Throughput Library Prep kit (Illumina, 20015963) with TruSeq RNA CD Indexes (Illumina, 20015949). Libraries were sequenced using Illumina NextSeq500, generating 20 million reads (75 bp, paired-end sequencing, whole cortex). INTACT-purified nuclear RNA (Norgen, 51800) was converted to cDNA and amplified with the Ovation RNA-seq System V2 (Nugen, 7102-A01) and libraries prepared with Kapa LTP library preparation kit (Illumina, KK8232) with TruSeq DNA Single Indexes (Illumina, 20015960), generating 40 million reads (150 bp, paired-end sequencing using Illumina HiSeq 4000 or NovaSeq 6000).

### Whole-genome bisulfite sequencing

DNA methylome libraries were generated using a modified snmC-seq method adapted for bulk DNA samples as previously described.^71^ 1% unmethylated lambda DNA (Promega, D1521) was added to each sample. Libraries were sequenced using an Illumina HiSeq 4000 instrument. The mapping of DNA methylome reads was performed as described.^33,71^

### Whole-cell patch clamp recordings

Male C57Bl/6J mice (P6-P17) were anesthetized with isoflurane and decapitated. The brains were quickly removed and coronal slices of the frontal cortex containing the prelimbic region (∼2 mm anterior to Bregma) were cut in an ice-cold slicing medium of the following composition (in mM): 110 sucrose, 2.5 KCl, 0.5 CaCl_2_, 7 MgCl_2_, 25 NaHCO_3_, 1.25 NaH_2_PO4, and 10 glucose (bubbled with 95% O2 and 5% CO_2_). The slices were then transferred to artificial CSF (aCSF) containing (in mM): 130 NaCl, 2.5 KCl, 1.25 NaH_2_PO4, 23 NaHCO_3_, 1.3 MgCl_2_, 2 CaCl_2_, and 10 glucose, equilibrated with 95% O_2_ and 5% CO_2_ at 35°C for 30 min and afterward maintained at room temperature (22–24°C) for at least 1 hr before use. Brain slices were then transferred to a recording chamber and kept minimally submerged under continuous superfusion with aCSF at a flow rate of ∼2 ml/min. Whole-cell patch-clamp recordings were obtained from putative L2/3 and L5 pyramidal neurons in the prelimbic cortex (PL), identified by their pyramidal-shaped cell bodies and apical dendrites extending toward Layer 1, as observed using an upright microscope equipped with differential interference contrast optics. Pipettes had a tip resistance of 4–8 MΩ when filled with an internal solution of the following composition (in mM): 125 K-gluconate, 15 KCl, 8 NaCl, 10 HEPES, 2 EGTA, 10 Na_2_ phosphocreatine, 4 MgATP, 0.3 NaGTP (pH 7.25 adjusted with KOH, 290–300 mOsm). Access resistance (typically 15–35 MΩ) was monitored throughout the experiment to ensure stable recordings.

After obtaining the whole-cell configuration in voltage-clamp mode, cells were switched from a holding potential of –70 mV to current-clamp mode, and the bridge-balance adjustment was performed. Membrane resistance was quantified from recordings with hyperpolarizing current injections that evoked small ∼5 mV deflections in membrane potential from resting. Stepwise current injections (0 pA to determine V_rest_, then increasing in 10 pA increments for 1 s) were recorded at 20 kHz to calculate rheobase - the minimum current required to elicit the first action potential. Action potential properties were measured from this initial spike. Input-output (I-O) curves were then constructed from the same recordings to assess neuronal firing responses across a range of depolarizing currents. Cells were then switched back to voltage-clamp mode (V_h_=−70 mV), and miniature excitatory postsynaptic currents (mEPSCs) were recorded in the presence of Na^+^ channel blocker, TTX (0.5 μM) and picrotoxin (50 μM) to prevent the generation of action potentials and minimize inhibitory responses. In these conditions, mEPSCs could be blocked by the AMPA receptor antagonist, CNQX (25 μM). Single events larger than 6 pA were detected offline using the Minianalysis program (Synaptosoft Inc Decatur, GA). All data were acquired using a Multiclamp 700B amplifier and pCLAMP 9 software (Molecular Devices, LLC, San Jose, CA). Data were analyzed using a two-way ANOVA with age and treatment as factors, followed by Sidak’s multiple comparisons test. Input-output (I-O) curves were analyzed using repeated-measures ANOVA, with stimulus intensity (10 pA increments) as the repeated factor, and treatment as the group factor. This analysis tested whether firing responses differed between treatment groups across stimulus intensities. Statistical significance was set at p<0.05.

### Sample selection procedure

Our experimental design included sampling time points that align critical developmental stages: E15, P0, and postnatal weeks 1 and 2. Although our electrophysiological analyses (P6-17) follow shortly after the epigenomic profiling at birth (P0), this timeline aligns with critical developmental windows where early epigenetic disruptions can manifest physiologically within days. Future studies with more closely spaced sampling could further refine this temporal resolution.

During the course of our experiments, we observed high inter- and intra-litter variability in transcriptional alterations after PIC-MIA (Supplementary Fig. 4). To be able to determine the offspring that should undergo transcriptional and methylome analyses, we conducted a study of the dam’s weight change after PIC injection. The effectiveness of PIC-MIA was confirmed by the reduction in the pregnant dams’ daily weight gain following injection at E12.5^72,73^ (Supplementary Fig. 4a). We observed that analysis of the dam’s daily weight changes (weight of E13.5 - weight of E12.5)/weight of plug day) showed a marked change in weight gain during the 24-48h post-treatment. Computational modeling using conditional inference tree-which tests the global null hypothesis of independence between any of the input variables and the response, and selects the time point that best discriminates the condition-suggested that a minimum of 5% change in weight at E13.5 was a good predictor of treatment response for both INTACT and wild-type mice. Therefore, the offspring of PIC-MIA dams were selected based on the reduction in the pregnant dams’ daily weight gain following injection at E12.5. Control (saline-injected) animals were excluded (20% of total samples) if their weight gain was less than 5% of that at E13.5, while PIC-MIA animals were excluded (17% of total samples) were excluded if their weight gain exceeded 5% of that at E13.5.

To select those pups that would undergo methylome analysis, we selected the most representative samples in each group defined by the gene expression profile as follows: First, using our transcriptome datasets from pyramidal nuclei, we performed dimensionality reduction to extract the global patterns of gene expression based on principal component (PC) analysis (Supplementary Fig. 2b,c). Second, given PC scores, we used linear discriminant analysis to train a classifier to distinguish PIC- and SAL-treated samples. Using this classifier, we assign a score to each sample that indicates the strength of the treatment effect on a per-animal basis. Finally, we selected 6 samples from each treatment group (PIC and SAL) which had the largest score based on our classifier. We have a representation of 3 litters for each treatment condition that underwent methylation sequencing and analysis.

### Transcriptome data analysis methods

Bulk RNA-seq read quality was assessed using FastQC.^74^ STAR (2.5.1b)^75^ was used to map sequence reads to the mouse mm10 reference genome and RSEM (version 1.2.31)^76^ was used to quantify gene expression. All samples were processed concurrently within each time point, eliminating batch effects at E15 and P13 (Supplementary Fig. 2b, metadata). However, P0 samples were run in multiple cohorts, resulting in strong batch effects (Supplementary Fig. 2c). To mitigate batch effects observed in RNA-seq data at P0, we applied ComBat-seq correction.^77^ PCA analyses before and after correction (see Supplementary Fig. 2d) demonstrate the successful normalization of batch effects, ensuring robust downstream transcriptional analyses. Mitochondrial and lowly expressed genes with fewer than 10 raw counts in all but one sample were removed before differential gene expression analysis. Differential expression analysis was conducted using DESeq2^78^ with a false discovery rate 0.1. Enrichment analysis was performed by inputting differentially expressed genes into EnrichR.^37,38^ Excitatory cortical genes were identified using Molecular Signatures Database^35^ mouse gene sets.^36^

### Differentially methylated region (DMR) analysis

After selecting the 12 samples from the PIC and SAL groups, we employed dispersion shrinkage for sequencing data (DSS) functions (v2.48.0),^79^ DMLtest, and callDMR, to detect DMRs between the two treatment groups. First, we extracted information from each sample regarding the characteristics of each CpG site within each chromosome. This includes the chromosome number, genomic coordinate, total cytosine base count, and the count of methylcytosines. We transformed the resulting dataset into a BSSeq file, which served as the input for the DMLtest function. Second, we use the DMLtest function to find significant differentially methylated loci (DML). DMLtest assumes that the data has a beta-binomial distribution, and estimates mean methylation levels and dispersions for all CpG sites. To improve estimation accuracy, methylation levels at each CpG site were smoothed by incorporating information from neighboring sites. Following this, the function conducted Wald tests at each CpG site to determine a test statistic for the significance of the difference in mean methylation between the two treatment groups. Third, we use the callDMR function to merge the resulting DMLs into DMRs. Specifically, this function selected DMLs with p-values below 1e-5 in the Wald test and combined nearby loci to form DMRs. A minimum DMR length of 50 base pairs and a minimum of 3 CpG sites were set as criteria. To form a DMR, CpG sites must be within a maximum distance of 50 base pairs from each other. Additionally, for each DMR, more than 50% of the CpG sites within this region must be significant. Fourth, we categorized the DMRs into hypo-DMRs and hyper-DMRs based on the difference in methylation levels between the PIC and SAL groups. Hypo-DMRs are defined as regions where the difference in methylation levels between the PIC and SAL groups is less than −0.1, and Hyper-DMRs are defined as regions where the difference is greater than 0.1.

### Enrichment Analysis of MIA DMRs and DEGs

In order to understand the regulatory dynamics between the PIC-MIA DMRs and DEGs, we performed an enrichment analysis to identify the DEGs at P0 that are associated with DMRs at P0. We split the DEGs into up- and down-regulated groups. Up-regulated DEGs are the genes that have higher expression levels in the PIC-MIA group than in the control group, and down-regulated DEGs are the genes that have lower expression levels in the MIA group than in the control group. We also split the DMRs into hypo- and hyper-methylated groups. Hypo-methylated DMRs are the RNA regions that are less methylated in the PIC-MIA group than in the control group, and hyper-methylated DMRs are the RNA regions that are more methylated in the PIC-MIA group than in the control group. We then used bedtools utilities (v2.31.0)^80^ to identify the overlaps between hyper- and hypo-methylated DMRs with up- and down-regulated DEGs. To account for the DMRs that do not directly overlap with DEGs but are adjacent to some DEGs and thus could potentially regulate them, we extended the DEGs by 5000 base pairs upstream and 5000 base pairs downstream using bedtools slop.

We performed two enrichment analyses: the enrichment of hypo- and hyper-methylated PIC-MIA DMRs in up-regulated DEGs, and the enrichment of hypo- and hyper-methylated PIC-MIA DMRs in down-regulated DEGs. We quantified the extent of overlaps by counting the number of overlapping regions in each intersect file. To determine whether the overlaps occurred by chance or reflected a true association, we conducted a bootstrapping analysis. We shuffled the hypo- and hyper-methylated DMRs multiple times and recalculated their overlaps with the DEGs. If the overlap with hypo- or hyper-methylated DMRs significantly decreased after shuffling, it would suggest a real association between this type of DMRs and the DEGs. We combined all hypo- and hyper-methylated DMRs, shuffled them, and then randomly reassigned them, keeping the number of DMRs for each type constant. We then located the intersections and measured the extent of overlaps in a manner identical to that of the unshuffled intersections. We repeated this process 500 times, generating 500 sets of overlap measurements.

To assess the differences between shuffled and unshuffled DMRs, we applied two methods: the observed/expected (unshuffled/shuffled) ratio and the p-value of unshuffled overlap within the distribution of the shuffled overlaps. To calculate the observed/expected ratio, we calculated the average overlap for hypo- and hyper-methylated DMRs over the 500 sets and divided the unshuffled overlap by it. To obtain the p-values, we counted how many simulated (shuffled) values were as extreme as or more extreme than the observed (unshuffled) value and calculated the empirical p-value as the proportion of extreme counts over the total number of shuffles. We defined MIA DMRs as enriched in a particular cell type if they met the following criteria: an observed/expected ratio greater than 1.1 and a p-value of less than 0.01.

### Enrichment Analysis of Hypo and Hyper MIA DMRs in Association with Cell-Type DMRs

To understand whether cell-type-specific DMRs were affected by MIA, we performed enrichment analysis between hypo- and hyper-methylated MIA DMRs and cell-type DMRs. To identify the overlap between MIA DMRs and cell type-specificDMRs, we conducted six enrichment analyses with three previously published single-cell DNA methylome datasets (snmC-seq), using bedtools utilities (v2.31.0).^80^ We first used the intersect function to determine the overlaps between the MIA DMRs (both Hyper-DMRs and Hypo-DMRs) and the DMRs from previously identified mouse frontal cortex cell types.^29^ We also calculated the overlaps between the MIA DMRs and the DMRs from previously identified human frontal cortex neuron types.^29^ To compare the MIA DMRS and DMRs from human genome coordinates, we used the liftOver utility^81^ to transform the MIA DMRs to hg19 genome coordinates. Finally, we calculated the overlaps between the MIA DMRs and the DMRs from previously identified mouse primary motor cortex cell types.^46^

For each of these six enrichment analyses, we quantified the extent of overlaps by counting the number of regions in each intersect file. To verify whether the overlaps occurred by chance or were due to an actual association, we used bootstrapping analysis. We shuffled the DMRs between different cell types and recalculated the overlaps. If the overlap with a particular cell type significantly decreased after shuffling, it would suggest a real association between the MIA DMRs and the DMRs of that cell type. We combined all cell-type-specific DMRs, shuffled them, and then randomly reassigned them back to the respective cell types, keeping the number of DMRs for each cell type constant. We then located the intersections and measured the extent of overlaps, in a manner identical to that of the unshuffled intersections. We repeated this process 500 times, generating 500 sets of overlap measurements. To assess the differences between shuffled and unshuffled DMRs, we applied two methods: the observed/expected (unshuffled/shuffled) ratio and the p-value of unshuffled overlap within the distribution of the shuffled overlaps. To calculate the observed/expected ratio, we calculated the average overlap for each cell type over the 500 sets and divided the unshuffled overlap by it. Next, to obtain the initial p-values, we calculated how many standard deviations the observed data point was from the mean of its simulated distribution for each cell type using the Z-statistic. These calculated p-values were subsequently ranked in ascending order. We then adjusted these p-values following the Benjamini-Hochberg procedure, where each p-value was adjusted as: Adjusted P-value = (Initial P-value × Total Number of Comparisons) / Rank. This approach controls the False Discovery Rate, which is crucial in multiple hypothesis testing scenarios, as it helps reduce the likelihood of false positives, thereby ensuring the statistical robustness of our findings. We defined MIA DMRs as enriched in a particular cell type if they met the following criteria: an observed/expected ratio greater than 2 and an adjusted p-value of less than 0.01.

### Enrichment Analysis of Overlapping MIA and Cell-Type DMRs in Association with DEGs

Recent single-nucleus analysis has shown that DNA methylation patterns can distinguish cell types in the brain.^24,29,30^ To gain insight into whether the DMRs affected by PIC-MIA overlap with cell-type-specific DMRs and how this may influence differential gene expression, we performed enrichment analyses between the PIC-MIA-affected cell-type-specific DMRs and the PIC-MIA DEGs. The PIC-MIA-affected cell-type DMRs were defined as the overlaps between a certain cell type’s DMRs and the PIC-MIA DMRs. We identified PIC-MIA-affected cell-type DMRs by using previously identified mouse frontal cortex cell types^29^ and the DMRs from previously identified mouse motor cortex cell types.^46^ We then identified their overlaps with both the up-regulated and down-regulated PIC-MIA DEGs at P0. We applied bootstrapping analysis to test the significance of the association between the MIA-affected cell-type DMRs and the DEGs. We first calculated the unshuffled overlaps between each MIA-affected cell-type DMR and the DEGs. We then shuffled all MIA-affected cell-type DMRs 500 times and calculated the overlaps for each shuffle. We calculated the observed/expected ratios and adjusted p-values identical to the enrichment analysis described above. Significant associations between the MIA-affected cell-type DMRs and the DEGs were defined as having an observed/expected ratio larger than 2 and an adjusted p-value less than 0.15.

## Supporting information

Sample Metadata

## Data access

All sequencing data are available in the Gene Expression Omnibus under accession GSE293592.

To review GEO accession GSE293592:

Go to

https://urldefense.com/v3/__https://www.ncbi.nlm.nih.gov/geo/query/acc.cgi?acc=GSE293592__;!!GX6Nv3_Pjr8b-17qtCok029Ok438DqXQ!yuPWcK0w41mNUqgLszrP6AF5DuKD5W8y_-jFd3r4xShKgpUTSog5qqUsKH2EEpO6zSAR9BNphmtdI5cg$

Enter token wzqhwsewnhmtzil into the box

## Acknowledgments

We thank Dr. Carolyn O’Connor, Director of the Flow Cytometry Core at Salk, for her input on the purification of nuclei. We are grateful to Joseph Chambers for animal treatment and care. This work was supported by grants from NIH: ES025585 to MMB, SBP, and JRE; MH112763 to MMB and JRE. HHMI to JRE. J.A. was a recipient of a UC San Diego Fellowship. The Flow Cytometry Core Facility of the Salk Institute (RRID:SCR_014839) is supported by funding from NIH-NCI CCSG P30 CA014195, and Shared Instrumentation Grants S10-OD023689, and S10 OD034268. J.R.E. is an investigator of the Howard Hughes Medical Institute.

## Author contributions

CYL: designed and conducted experiments, analyzed data, manuscript writing

JA: analyzed data, manuscript writing

APD: designed and conducted experiments, analyzed data, manuscript writing

SW: analyzed data

JL: analyzed data, manuscript editing

HL: analyzed data, manuscript editing

JO: conducted experiments

RGC: conducted experiments

JN: conducted experiments

SBP: Conceptualization, design and supervision, funding acquisition, review and editing

JRE: Conceptualization, design and supervision, funding acquisition, review and editing

EAM: Conceptualization, design and supervision, funding acquisition, review and editing

MMB: Conceptualization, design and supervision, funding acquisition, review and editing

---

**Supplementary Fig. 1.**
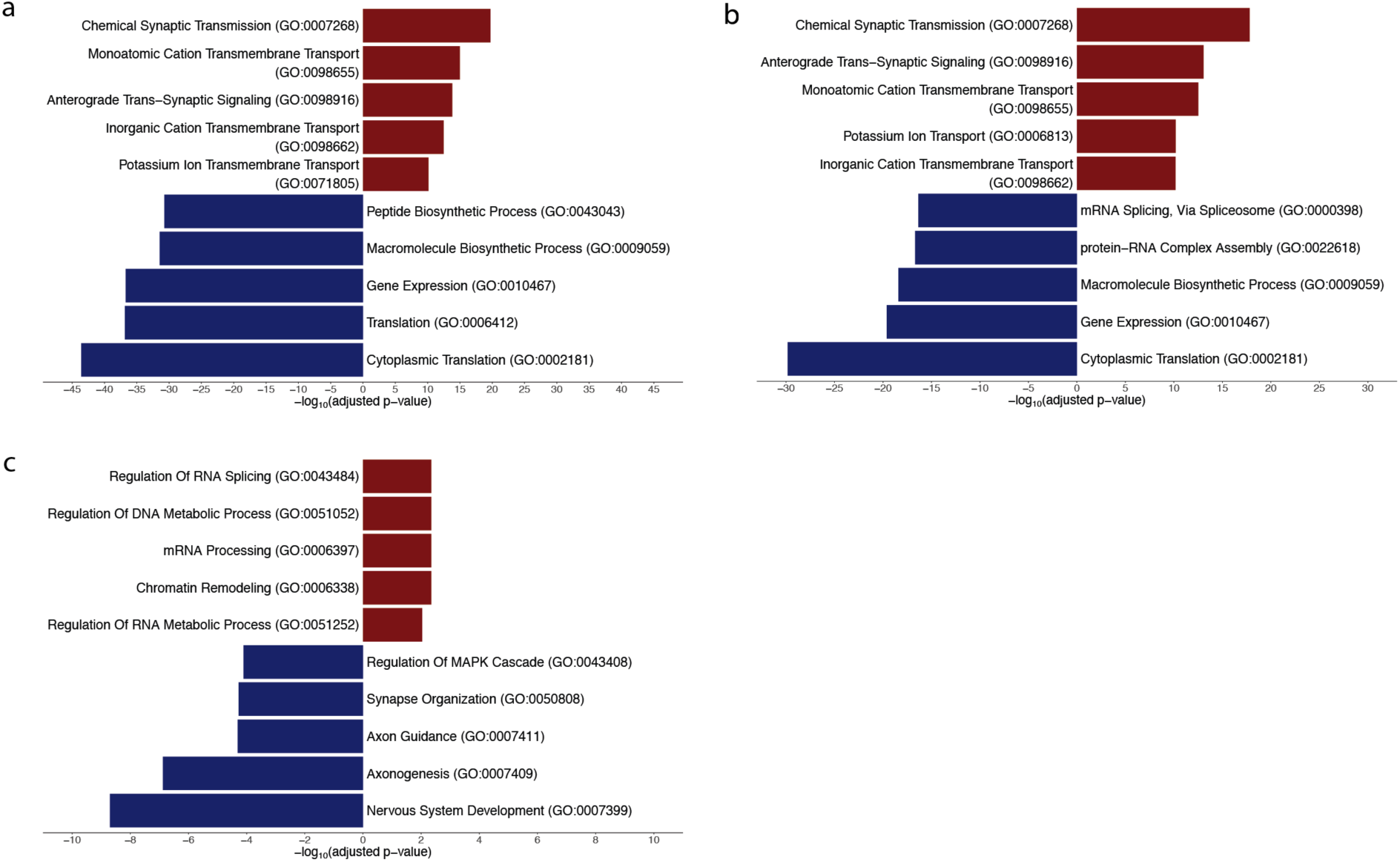
Gene Ontology Biological Process Enriched Terms in Normal Development. All categories have significant adjusted p-values (FDR<0.05). **a.** Enriched terms between P0 and E15 DE protein-coding genes. Bars pointing to the left are enriched at E15 (colored in blue) and to the right are enriched at P0 (colored in red). **b.** Enriched terms between P13 and P0 DE protein-coding genes. Bars pointing to the left are enriched at P0 (colored in blue) and to the right are enriched at P13 (colored in red). **c.** Enriched terms between P39 and P13 DE protein-coding genes. Bars pointing to the left are enriched at P13 (colored in blue) and to the right are enriched at P39 (colored in red).

**Supplementary Fig. 2.**
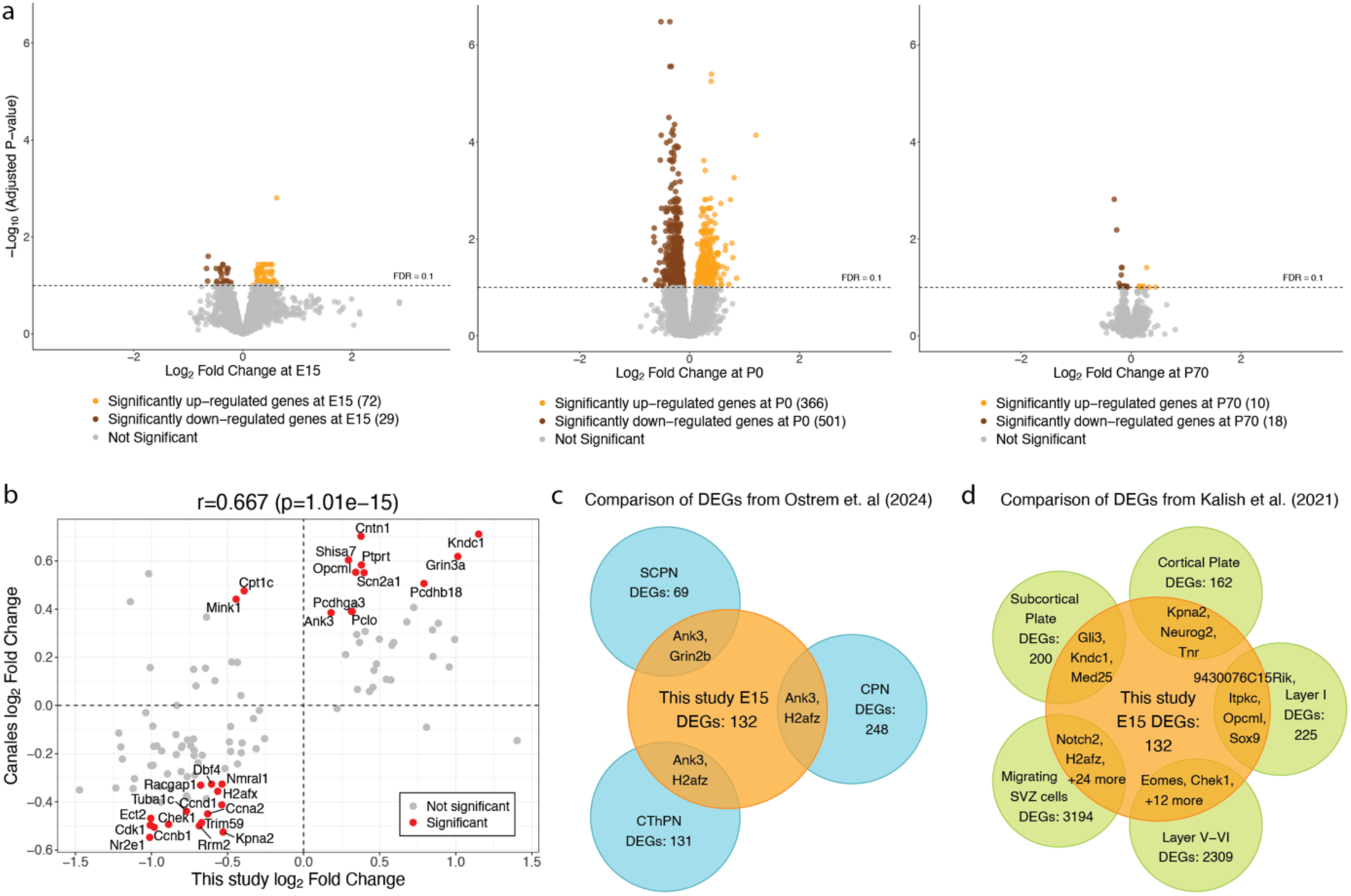
Global transcriptional changes in the perinatal cortex following PIC-MIA. **a.** Volcano plots with a log fold change of gene expression against adjusted p-value showing the transcriptional changes in the cortex of offspring of PIC vs. SAL at E15 (left), P0 (center), and P70 (right). Yellow points are up-regulated and brown are down-regulated genes in PIC. Grey dots are not significantly differentially expressed genes. **b.** Comparison of DE genes at E15 identified in our excitatory cortical neuron data and in Canales et al.^22^ where points in red are significant in both datasets. **c.** Overlap of PIC-MIA DE genes identified in our E15 excitatory cortical neuron data (yellow) and in somatosensory cortical neuron populations from single-nuclei transcriptomes^43^ (blue). CPN, callosal projection neurons; SCPN, subcerebral projection neurons; CThPN, corticothalamic projection neurons. **d.** Overlap of DE genes identified in our E15 excitatory cortical neuron data and in single-cell RNA-seq at E14^42^ (FDR<0.1). SVZ, subventricular zone.

**Supplementary Fig. 3.**
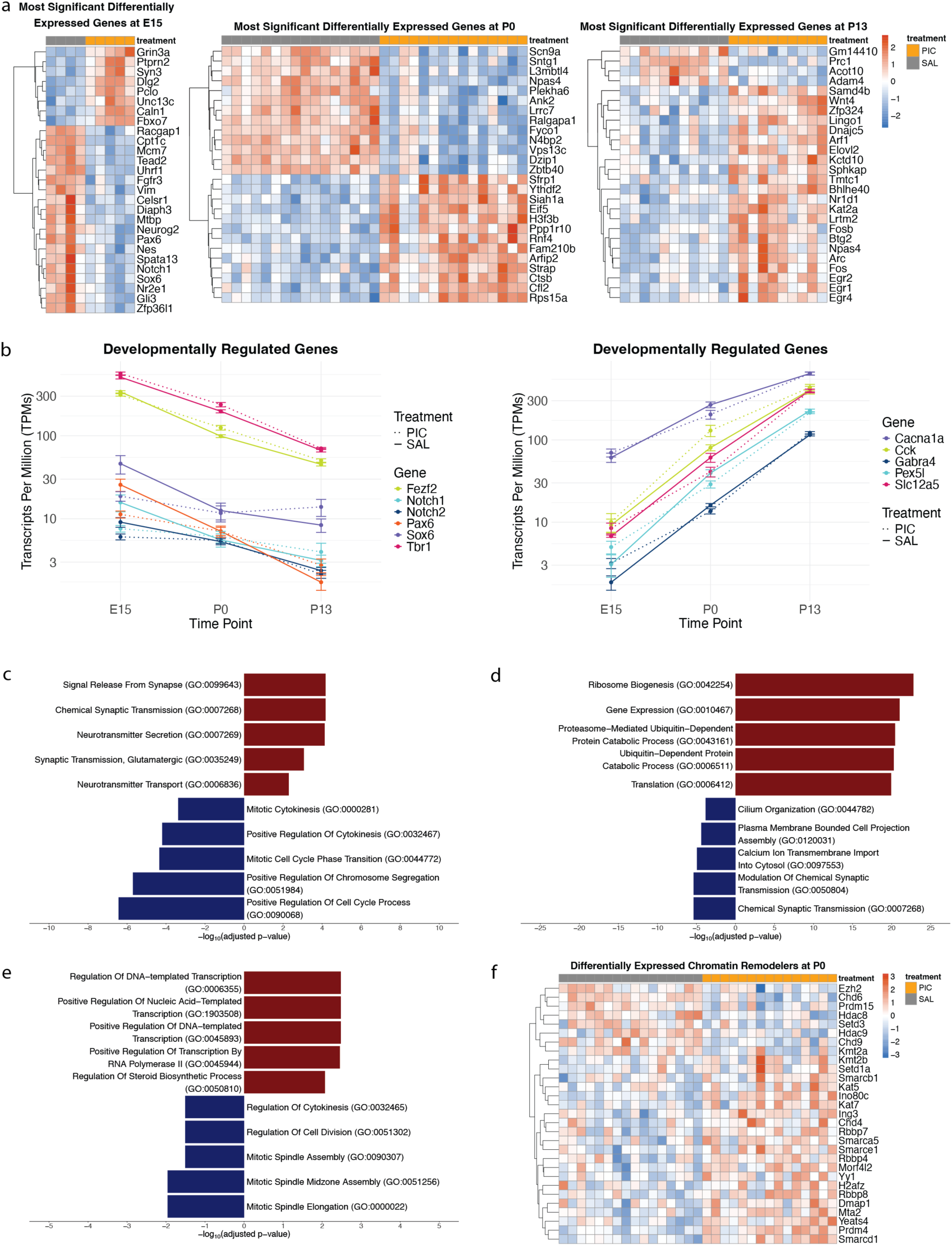
PIC-MIA effects on gene expression of excitatory cortical neurons. **a.** Distribution of the top DE genes at E15 (left), P0 (center), and P13 (right). Heatmaps were plotted using variance-stabilized expression values, with each gene re-scaled by Z-score normalization and ordered by hierarchical clustering. The red color indicates higher expression levels while the blue denotes lower expression. **b.** Trends in averaged TPMs for genes that go up (left) and down (right) with development. c-e. Gene Ontology Biological Process Enriched Terms in protein-coding DE genes at E15 (**c**), P0 (**d**), and P13 (**e**). Bars pointing to the left are enriched in SAL (colored in blue) and to the right are enriched in PIC (colored in red). All categories have significant adjusted p-values (FDR<0.05). **f.** Distribution of DE genes at P0 involved in chromatin remodeling.

**Supplementary Fig. 4.**
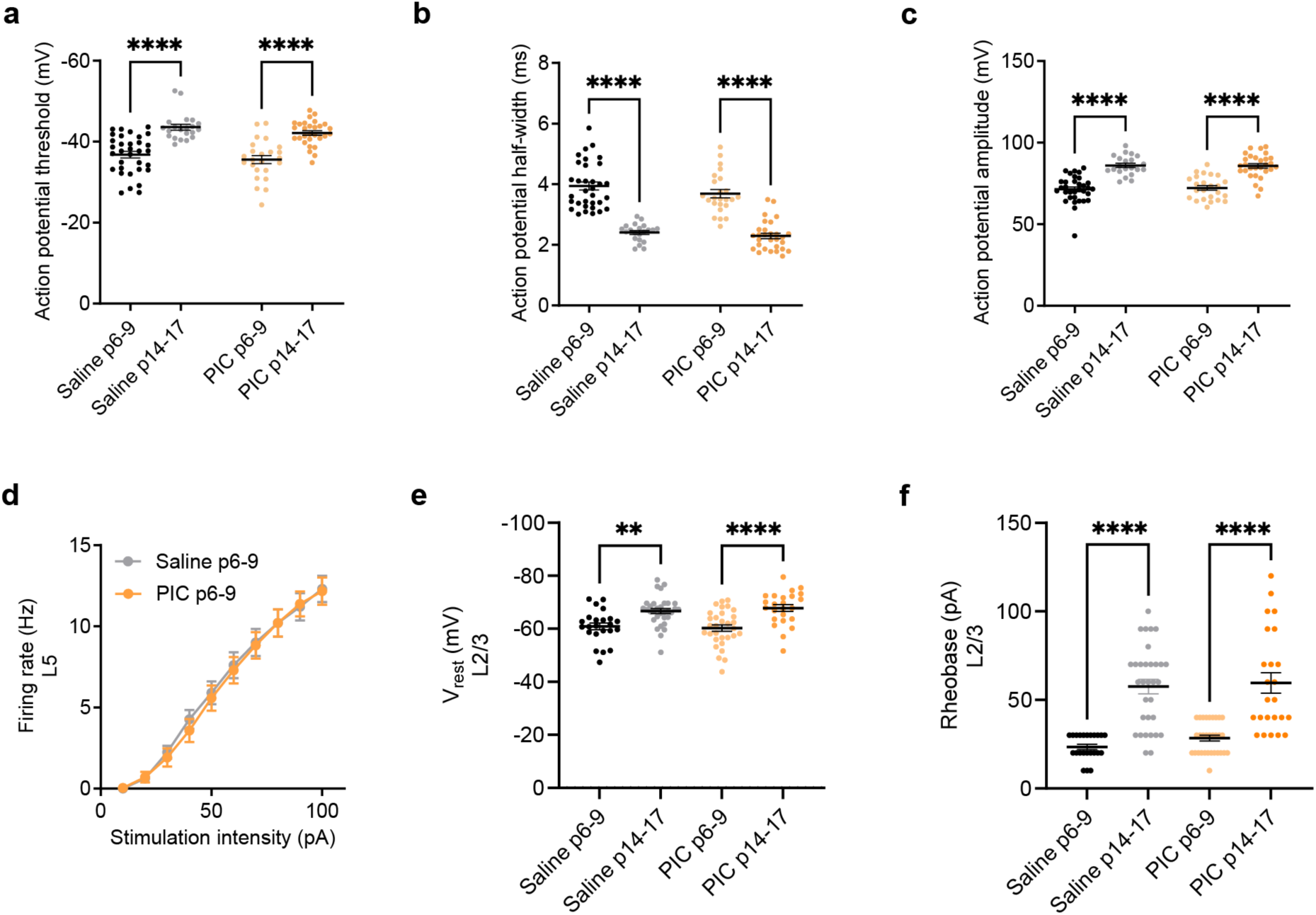
Developmental changes unaffected by PIC treatment. Comparison of saline- and PIC-treated groups at two developmental time points (P6–9 and P14–17) across several properties of L5 (a–d) and L2/3 pyramidal neurons (e–f). **a.** Action potential threshold became more negative over time, and action potential half-width shortened (**b**). **c.** Action potential amplitude increased developmentally in both groups. **d.** Input-output curves showed no significant differences between groups at P6–9. L2/3 pyramidal neurons exhibited developmental hyperpolarization (**e**), and rheobase increased (**f**) over time in both conditions. Data points represent individual neurons, with mean values indicated by horizontal bars. Error bars denote SEM. Statistical significance: **p<0.01, ****p<0.0001.

**Supplementary Fig. 5.**
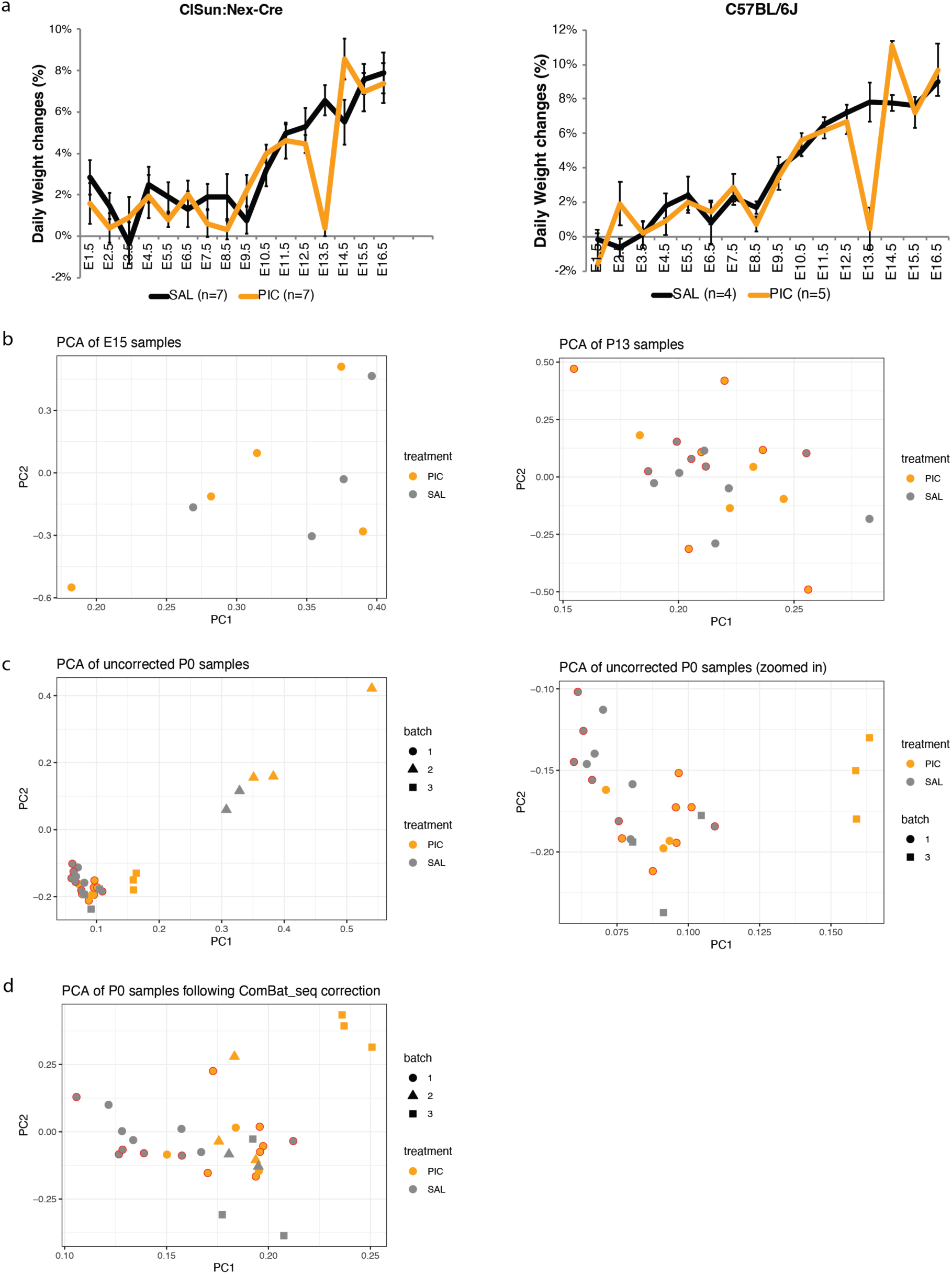
MIA Sample Pre-Processing. **a.** Daily Weight changes of pregnant dams after Poly(I:C) (PIC) or saline (SAL) treatment. Pregnant dams were weighted each day at the same time of day. A clear drop (p = 1 × 10-5) in weight gain is observed one day after injection (E13.5) in the PIC group of either Clsun: NexCre (left) or C57BL/6J mice(right). Mean and standard error were plotted. Daily weight change = (weight of E13.5 - weight of E12.5)/weight of plug day. **b**. Principal components of RNA-seq of time points with no batch effects. **c.** PCA of P0 samples with batch effects. d. PCA of P0 samples after ComBat_seq batch correction.

